# Phage-phage competition and biofilms reduce the efficacy of a combination of two virulent bacteriophages against *Pseudomonas aeruginosa*

**DOI:** 10.1101/2024.09.13.612609

**Authors:** Magdalena Bürkle, Imke H. E. Korf, Anne Lippegaus, Sebastian Krautwurst, Christine Rohde, Chantal Weissfuss, Geraldine Nouailles, Xavière Menatong Tene, Baptiste Gaborieau, Jean-Marc Ghigo, Jean-Damien Ricard, Andreas C. Hocke, Kai Papenfort, Laurent Debarbieux, Martin Witzenrath, Sandra-Maria Wienhold, Gopinath Krishnamoorthy

## Abstract

Combined use of virulent bacteriophages (phages) and antibiotics reduces the severity of difficult-to-treat *Pseudomonas aeruginosa* infections in many patients. *In vitro* methods that attempt to reproduce more than one physiological state of the pathogens can provide a valuable assessment of antibacterials like phages. Here, by measuring bacterial killing kinetics and individual replication in different growth conditions, including biofilms and a human lung epithelial cell line, we elucidated factors influencing the efficacy of two virulent phages against *P. aeruginosa* PAO1. A single administration of phages effectively reduced the *P. aeruginosa* viability in planktonic conditions and infected human lung cell cultures, however, the emergence of phage-resistant variants occurred subsequently. In static biofilms, the phage combination displayed initial inhibition of biofilm dispersal, but sustained control was achieved only by combining phages and meropenem. In contrast, surface-attached biofilms exhibited tolerance to phage and/or meropenem, suggesting a spatiotemporal variation in the antibacterial effect. Moreover, the phage with the shorter lysis time lysed *P. aeruginosa* more rapidly and selected a specific nucleotide polymorphism that conferred a competitive disadvantage and cross-resistance to the second phage of the combination. The sequential addition of phages resulted in their unimpeded replication with no increase in bacteriolytic activity. These findings highlight biofilm developmental stages, phage-phage competition, and phage resistance as factors restricting the *in vitro* efficacy of a two-phage combination. Our findings provide a framework for selecting and optimizing phage combinations for enhanced efficacy against *P. aeruginosa*, a metabolically flexible pathogen that undergoes specific adaptation within the infected lung.

## Introduction

*Pseudomonas aeruginosa* causes life-threatening infections, especially in patients with impaired lung function, chronic wounds and device-related infections [1]. Chronic infections select for *P. aeruginosa* variants that increasingly form biofilms – aggregates of bacterial cells embedded in an extracellular matrix – enabling them to withstand the action of antibacterial immunity and of many antibiotics [2, 3]. Furthermore, the widespread emergence of multidrug-resistant (MDR) strains and the slow progress in the development of new antibiotics have reinvigorated the search for novel adjunctive strategies for treating infections by MDR pathogens.

A virulent bacteriophage (or phage) is a virus that lyses bacterial hosts with high specificity. This offers the possibility of using such phages as an adjunct agent to the antibacterial action of antibiotics and host response for killing pathogens, including MDR strains [4–8]. However, one of the limitations is the narrow host range of phages, which varies even among genetically related strains of the same bacterial species. Combining multiple virulent phages for therapeutic application is therefore proposed to broaden the bacteriolytic activity against diverse clinical isolates and, supposedly, to evade anti-phage defence systems and diminish the development of phage-resistance [9].

The conventional soft-agar overlay plaque method enables rapid and semi-empirical selection of phage combinations. However, this method determines phage lytic effects only at a single time point and uses a synchronous bacterial culture that does not adequately represent key phenotypic features in the diseased host [10, 11]. For example, in chronically infected patients, *P. aeruginosa* switches between surface-attached and dispersed lifestyles contributing to spatiotemporal variations of gene expression at the population level, which overall could affect its susceptibility to phages [12]. Taking this into account, several studies have additionally used microtitre assays to identify phages with varied efficacy against static biofilms [13, 14]. In addition, the nature of interactions of virulent phages among themselves and/or with biofilms can be complex and is likely to vary with the bacterial genotype [15–19]. Obtaining detailed information on the effects of phage(s) targeting surface-attached or dispersal stages of a biofilm is essential for understanding how the specific *P. aeruginosa* growth adaptations within the biofilm structure affect the efficacy of phage combinations. In addition, the insights into the pattern of phage-phage interaction and correlations with antibacterial effects will increase the confidence in laboratory assessments of phage combination efficacy [20].

JG005 and JG024 are double-stranded (ds) virulent DNA phages belonging to the *Pakpunavirus* and *Pbunavirus* genera, respectively, and have been shown to lyse several *P. aeruginosa* isolates [21–24]. The primary objective of this study was therefore to identify the variables that influence the effects of the combination of JG005 and JG024 against planktonic and biofilm growth of *P. aeruginosa* PAO1. Based on bacterial killing kinetics and individual phage replication readouts, as well as dual RNA-Seq, and genome sequencing, our results have identified *P. aeruginosa* genotype, biofilm spatial restriction, phage resistance, and phage-phage competition as interrelated factors influencing the bacteriolytic efficacy of this two-phage combination.

## Materials and Methods

### Phages and bacterial strains

*P. aeruginosa* PAO1 (DSM 19880, Leibniz Institute DSMZ GmbH, Germany) and other strains (see Supplementary methods) were routinely cultured on BD^TM^ Columbia Agar with 5 % sheep blood or Luria-Bertani planktonic (LB, Carl Roth, 6669.2) with orbital shaking (200 rpm) at 37 °C. Bacterial growth was measured at 600 nm using a spectrophotometer (CO 8000 cell density meter). Single plaque of each phage stock (Leibniz Institute DSMZ GmbH, Germany) from LB agar plates were picked and propagated to produce chloroformed (1:1000 v/v) lysate and stored at 4 °C until further use. Chromatographically purified phage preparations (Fraunhofer Institute for Toxicology and Experimental Medicine, Germany) were used only for A549 infection experiments. Digital images of plates containing confluent plaques (Interscience Scan 4000) were transformed using FIJI ImageJ2 prior to Plaque Size Tool analysis [25]. Bacteriophage-insensitive mutants (BIMs) frequencies were determined as reported earlier [26].

### Planktonic culture

Indicated *P. aeruginosa* strains were cultured in LB broth under shaking conditions to OD_600_= 0.2-0.3 (37°C with 5% CO_2_) and diluted 100-fold prior to treatment with either JG005 or JG024 or a combination of both at a 1:1 phage-to-bacteria ratio. For sequential treatment, JG024 (1E+07 PFU/mL) was added first and JG005 (1E+07 PFU/mL) was added 2 hours later.

### A549 cell line infection

A549 human type II alveolar epithelial cells (American Type Culture Collection; CCL-185^TM^) that had undergone 8-15 passages were used. Cells were cultured in antibiotic-free Dulbecco’s Modified Eagle Medium containing 10% heat-inactivated fetal calf serum and 1% L-glutamine in a humidified chamber at 37 °C with 5% CO_2_. Cells were seeded the day before the infection into 12-well plates with a final cell density of 1E+05 cells/well. Equal numbers of PAO1 were used for cell infection. Chromatographically purified phages (provided by the Fraunhofer Institute for Toxicology and Experimental Medicine, Germany) were added simultaneously one hour after infection at two different phage-to-bacteria ratios (1:10 and 10:1) according to the bacterial burden of the initial infection. Bacterial and phage enumeration were performed 6 and 24 hours after phage treatment.

### Microtitre biofilm assay

Overnight grown culture of PAO1 or its Δ*retS* derivative was diluted (OD_600_= 0.1) and 200 µL was added into a round bottom microtitre plate (Sarstedt, micro test plate 96 well) and incubated at 37°C and 5% CO_2_ for 24 hours in a humid chamber. Subsequently, the liquid phase was carefully pipetted out and the surface-attached material was carefully washed three times with sterile phosphate-buffered saline (PBS, Gibco™) and then treated with phage combinations, 5X MIC (5 µg/mL) meropenem (Sigma-Aldrich) or phage buffer alone. According to the initial bacterial viable counts, the phage combination (JG005, JG024, and Bhz17) was administered in two different phage-to-bacteria ratios (Phage_low_ 1:10, Phage_high_: 10:1, diluted in phage buffer). For sequential treatment, JG024 (5E+08 PFU in 50 µL) was added 2 hours prior to the addition of JG005 (5E+08 PFU in 50 µL).

The bacterial and phage counts were enumerated for the non-adherent and the adherent biofilm phases. First, the non-adherent population was carefully separated by pipetting and stored on ice for further processing. Second, the remaining adherent phase was washed three times with PBS (200 µL), followed by the addition of 100 µL of LB, and the biofilm was mechanically dislodged from the surface by vigorous scratching 30 times and pipetting up and down five times. The dislodging step was repeated with another 100 µL LB broth, and the suspension was collected. Samples were serially diluted 10-fold, plated on blood agar plates for bacterial counts, and spotted on a double agar overlay with each indicator strain for individual phage counts.

### Adsorption assay and One-step growth curve

One-step growth curve and adsorption rate of phage particles was determined under both mono- and coinfection conditions as previously described [22, 27]. For one-step curve, indicated phages and *P. aeruginosa* strains PAO1 or PA14 were incubated at a ratio of 0.01:1 for 10 minutes prior to sampling. Virion release at 50 minutes, since the phage titres plateaued there, was calculated as follows: (phage titre_50 minutes_ – phage titre_initial_)/(phage titre_initial_). The adsorption constant (K_m_) was calculated from the inverse value of the slope from a linear regression of the natural-log transformed free phages divided by the total amount of infected bacteria [28].

### JG005 phage genome

JG005 genome was annotated using CPT phage galaxy tools [29] followed by manual revision with UGENE [30] and HHPRED (https://toolkit.tuebingen.mpg.de/tools/hhpred). Default parameters were used for all software tools. Candidate tRNA or transfer-mRNA (tmRNA) genes were detected using ARAGORN [31] and tRNAscan-SE [32].

The bacterial RNA polymerase was inhibited with rifampicin (400 µg/mL, Sigma-Aldrich) to determine the dependency of the JG005, and JG024 on PAO1 RNA polymerases as previously described [33].

### Whole transcriptome sequencing and data analysis

Transcriptomic analysis via bulk RNA-Seq was carried out using logarithmic growth of PAO1, which was either mono-infected with phage JG024 and coinfected with phage JG024 and JG005 at phage-to-bacteria ratio 5:1to increase the chances of coinfection of the same single cell. Samples were collected as reported earlier with some modifications [27].

Total RNA was isolated by hot-phenol method and digested with TURBO^TM^ DNase (Thermo Fisher Scientific). Ribosomal RNA was depleted using rRNA-specific biotinylated probes [34]. cDNA libraries were prepared using the NEBNext® Small RNA Library Prep Set for Illumina (NEB; #E7300L). cDNA libraries were sequenced using a Illumina NextSeq1000 system in single-read mode for 100 cycles. Demultiplexed raw reads were trimmed for quality and 3’ adaptors using the CLC Genomics Workbench (Qiagen). Reads were mapped to PAO1, JG024 and JG005 genomic sequences (NCBI accession numbers: AE004091, NC_017674, PP712940) and unique gene reads mapped to the different organisms were compared to the total number of reads. Fold enrichment in the infected samples was compared to the untreated control using the CLC “Differential Expression for RNA-Seq” tool. Genes with a fold change ≥ 1.5 and an FDR-adjusted p-value ≤ 0.05 were defined as differentially expressed.

### Whole genome sequencing of phage-resistant mutants

Genomic DNA of *P. aeruginosa* phage resistant clones was extracted from overnight culture and sequenced by Illumina. Polymorphism were identified with the Snippy program (https://github.com/tseemann/snippy).

### Statistical Analysis

GraphPad Prism 10 software (San Diego, CA, USA) was used for statistical analyses. The specific statistical tests applied are indicated in the figure legends. *P* values of less than 0.05 were considered statistically significant.

### Data availability

Transcriptome sequencing data are available from the NCBI GEO database under accession number GSE271537.

## Results

### Phage treatment results in biphasic killing of *P. aeruginosa* in planktonic culture and in presence with alveolar epithelial human cells

At the outset, differences in plaque size and morphology were determined. JG005 and JG024 phages form clear plaques on the wild-type *P. aeruginosa* strain PAO1 lawn with mean sizes of 3.94 mm and 1.22 mm, respectively **(Fig. S1)**. We then evaluated the individual or combined activity of these two phages against a stirred planktonic culture of PAO1. The combined lytic effect of JG005 and JG024 was found similar to that of JG005. All phage treatments showed biphasic phenotypic spectrum: initial susceptibility followed by the emergence of phage-tolerant/-resistant bacterial subpopulation.

Then, we assessed the efficacy of the combination of JG005 and JG024 phages on PAO1 in presence with human lung epithelial cell line A549 monolayers at two different phage-to-bacteria ratios of: 1:10 (hereafter referred as phage_low_) or 10:1 (phage_high_) **(Fig. 1B)**. Phage treatments resulted in a similar biphasic killing effect in infected A549 monolayers comparable to that of planktonic culture conditions, with PAO1 growth strongly reduced at 6 hours and then resuming until 24 hours.

**Figure 1.**
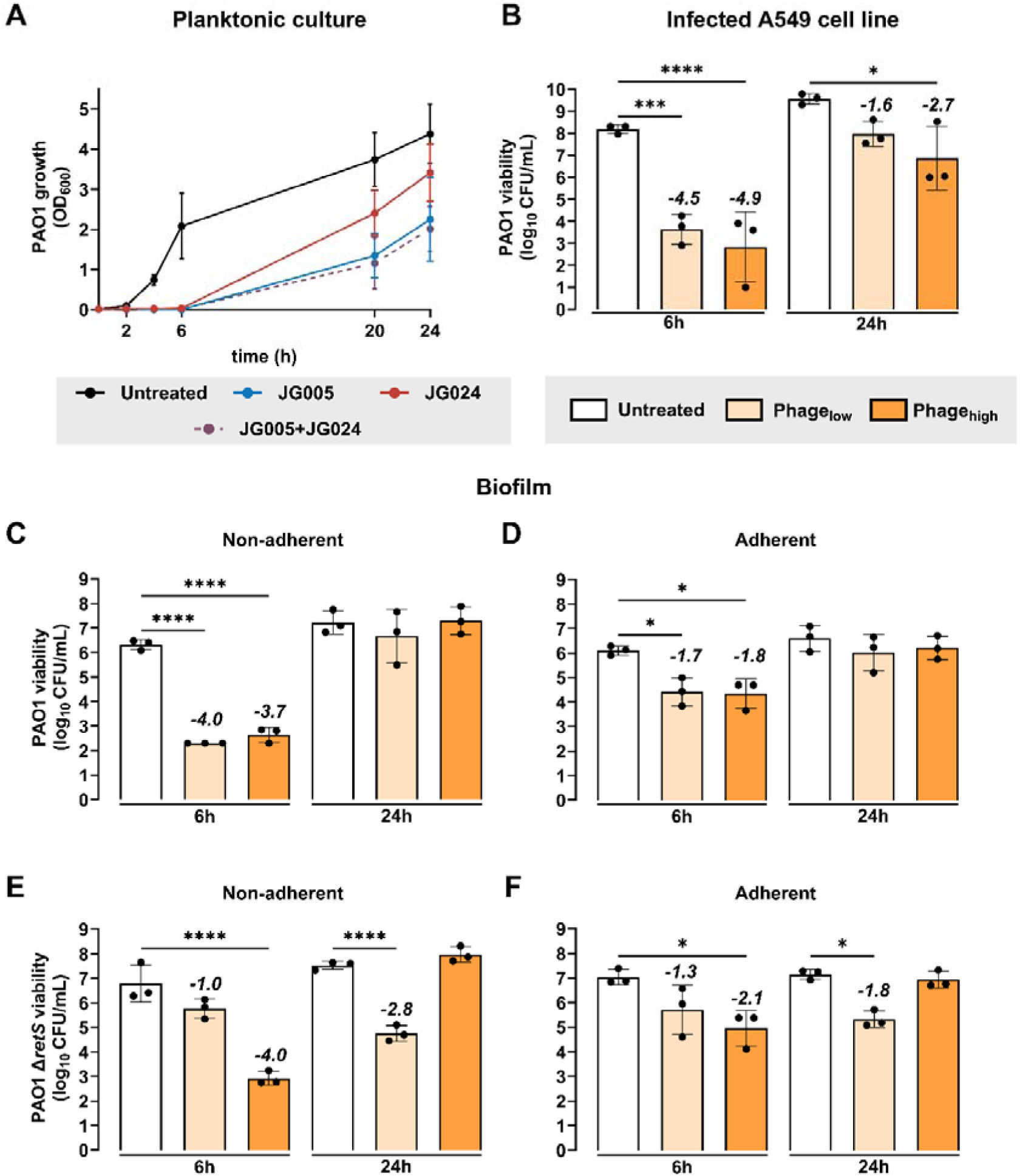
Time-kill-kinetics of phage combinations against different *P. aeruginosa* growth states. **(A)** *P. aeruginosa* PAO1 planktonic culture was treated with JG005, JG024, JG005+JG024 combination (phage-to-bacteria ratio 1:1) or with phage buffer as control. Bacterial growth was monitored by measuring optical density at 600 nm (OD_600_). **(B)** PAO1 viability from infected A549 human lung epithelial cell line at 6 and 24 hours post-treatment with single and combined treatment of JG005+JG024 at a phage-to-bacteria ratio of Phage_low_ with a 1:10, and phage_high_ with a 10:1 ratio, or untreated control. Phage treatment was performed one hour after cell line infection. Viability of PAO1 from **(C)** non-adherent (dispersed) and **(D)** adherent (surface-attached) biofilm phases, at 6 and 24 hours post-treatment with phage combination. Viability of PAO1 Δ*retS* **(E)** non-adherent and **(F)** adherent biofilm phases 6 and 24 hours post-treatment with phage combination. Data represent the mean ± SD of *n*= 3 biological replicates with technical duplicates. Numeric values on top of the columns indicate log-fold changes relative to the untreated control. Ordinary One-way ANOVA and Šídák’s multiple comparisons test was used to determine *P*-values (Not significant (*P* > 0.05) is not shown, * *P* < 0.05, *** *P* < 0.001, **** *P* < 0.0001). Detection limit is 200 CFU/mL, all samples below are set to this limit.

### Spatiotemporal constraints in biofilm reduce phage combination efficacy against *P. aeruginosa*

Next, we sought to establish the time-kill kinetics of phage combination against *P. aeruginosa* static biofilm. To this end, a combination of three phages (JG005+JG024+Bhz17) was added on a 24-hour-grown PAO1 biofilm in microtitre plates as schematically shown in **Fig. S2A**. Bhz17 is a dsDNA phage and cannot infect the PAO1 strain. Since its addition did not alter the time kill kinetics, Bhz17 is included as a control to monitor phage replication dynamics in biofilm **(Fig. S1A-B)**. Bacterial and individual phage counts were determined from two separated biofilm phases: surface-attached (adherent) and dispersed (non-adherent). After reduction of the viable PAO1 counts at 6 hours, a subpopulation grew to the level of the untreated control after 24 hours in both the adherent and the non-adherent phases, regardless of the differences in initial phage concentration used **(Fig. 1C-D).**

Interestingly, viable PAO1 counts in the non-adherent phase were lower **(Fig. 1C)** compared to the adherent phase at 6 hours after phage combination treatment **(Fig. 1D)**, indicating effective prevention of biofilm dispersion by phage treatment and little effect on the adherent population. No impact of the phage:bacterium ratio was noted in terms of bacterial viability reduction at this time point. These results reveal a spatiotemporal pattern in phage-mediated bacteriolysis, namely, a minimal impact on adherent population viability but an early inhibition of the biofilm dispersal.

We then assessed the lytic activity of the phage combination on biofilms with increased thickness, using a hyper biofilm-producing *P. aeruginosa* PAO1 Δ*retS* strain [35]. Biomass production, quantified by crystal violet staining, was found to be 2.4-fold higher in the untreated Δ*retS* biofilm compared to its wild-type **(Fig. S2B)**. Bacterial counts in the Δ*retS* biofilm were increased correspondingly **(Fig. 1C-F)**. The viability reduction kinetics of Δ*retS* biofilm upon phage_high_ combination treatment were comparable to those of wild-type strain at 6 and 24 hours post-treatment **(Fig. 1E-F)**. Counterintuitively, phage_low_ combination treatment reduced the Δ*retS* adherent and dispersed biofilm population over 24 hours, although at 6 hours the effect was weaker than in wild-type. Hence, the phage efficacy cannot be readily correlated to the increased biofilm formation of the *retS*-deficient strain. It is unclear whether this effect is due to a different phage stoichiometry resulting from the increased bacterial density or due to the RetS regulatory transcriptional network [36] itself.

### JG005 reduced JG024 replication in combination treatment

We also enumerated the titres of individual phages from each previous experimental condition. Counts of JG005 and JG024 showed an increase within two hours when applied individually in planktonic culture, and then plateaued. At 24 hours, JG005 titre (mean 1.36E+10 PFU/mL) was found to be higher relative to JG024 counts (mean 1.89E+09 PFU/mL) at 24 hours **(Fig. 2A, solid lines)**. In contrast, when combined **(Fig. 2A, dotted lines)**, only JG005 titre increased, closely matching the titres observed with individual treatment, while the titre of JG024 declined rapidly (mean 1.96E+05 PFU/mL) and then remained in a stalled status. Likewise, a decrease in JG024 titre with combined treatment was seen in the PAO1-infected A549 cell line **(Fig. 2B)**. Similar reduction in JG024 replicative fitness was also observed in PAO1 Δ*retS* and clinical *P. aeruginosa* strain CHA [37] planktonic cultures, although JG024 titre in the CHA strain reached the levels observed with single treatment by 24 hours **(Fig. S3A-B)**. To further investigate this phenomenon, *P. aeruginosa* PA14, not susceptible to JG005, was treated with a combination of JG005 and JG024. Notably, this resulted in uninhibited replication of JG024 (**Fig. S3C**), suggesting that the within-host replication of JG005 in PAO1 causes the reduced replication fitness of JG024.

**Figure 2.**
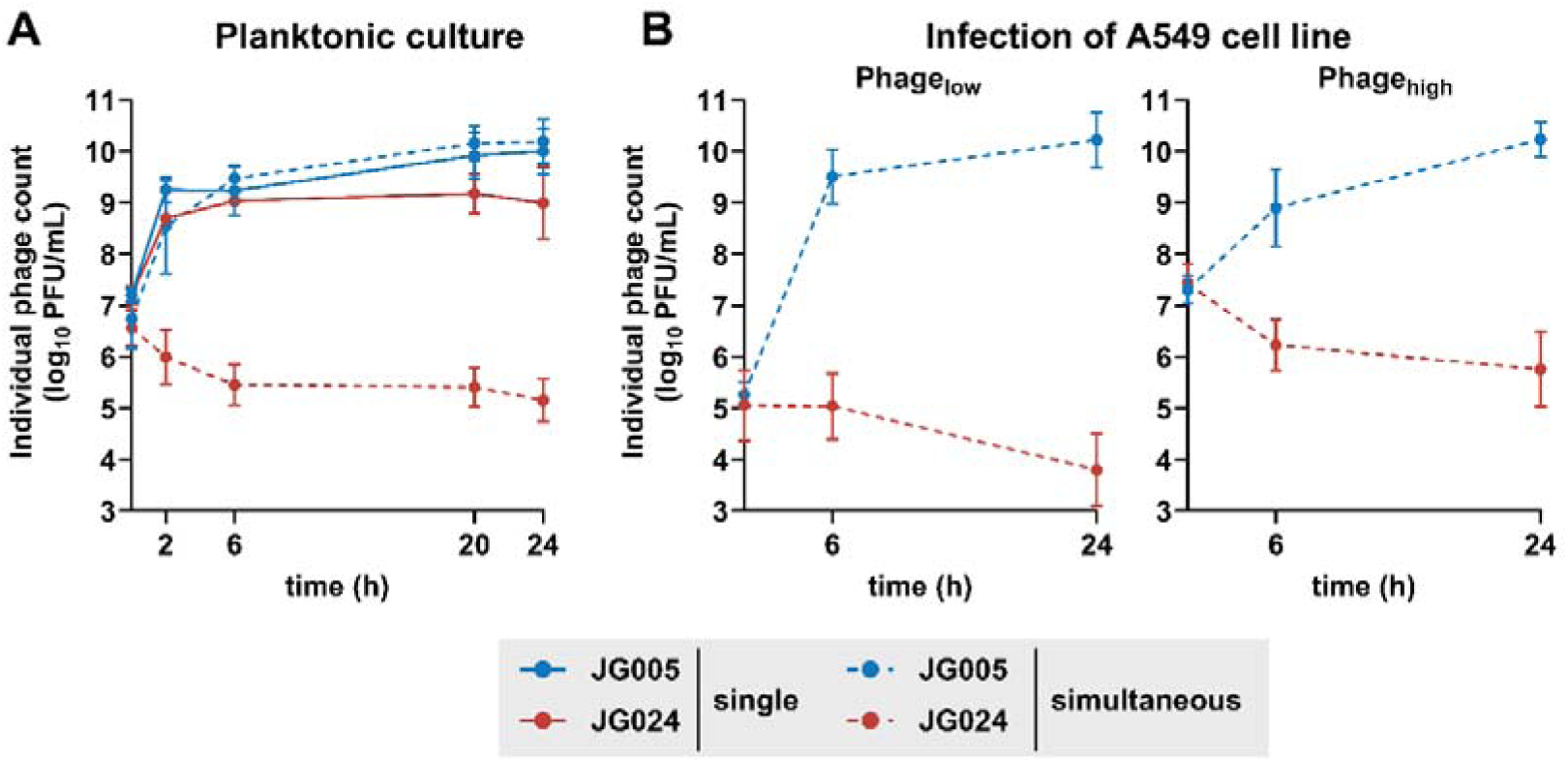
Individual phage replication kinetics of JG005 and JG024. **(A)** Both phages JG005 and JG024 were applied individually (single) or simultaneously at a phage-to-bacteria ratio of 1:1 to *P. aeruginosa* PAO1 planktonic culture or **(B)** in combination of at a phage-to-bacteria ratio of Phage_low_ with a 1:10, and phage_high_ with a 10:1 ratio to PAO1 in infected A549 human epithelial lung cell line monolayer. Data represent the mean ± SD of *n*=3 biological replicates. Individual phage titres were assessed by using specific *P. aeruginosa* indicator strains (**see supplementary methods**). Solid lines indicate single phage treatment, and dotted lines indicate simultaneous combined treatment.

### Phage abundance varies with biofilm phase

Phage titre in the separate phases of a PAO1 biofilm after combined treatment were also subsequently examined. JG005 titre increased in the non-adherent biofilm phase **(Fig. 3A),** albeit slower than in planktonic culture conditions and only at the phage_low_ concentration **(see Fig. 2A)**, which likely contributed to better inhibition of biofilm dispersal in the non-adherent phase **(see Fig. 1C)**. JG024 titre rapidly decreased by approximately two orders of magnitude particularly at the phage_high_ concentration, similarly as what we observed in planktonic condition, and were occasionally even lower than titres of the PAO1-non-replicative Bhz17. The phage titres in the adherent biofilm showed a similar pattern but dropped even lower than in the non-adherent phase.

**Figure 3.**
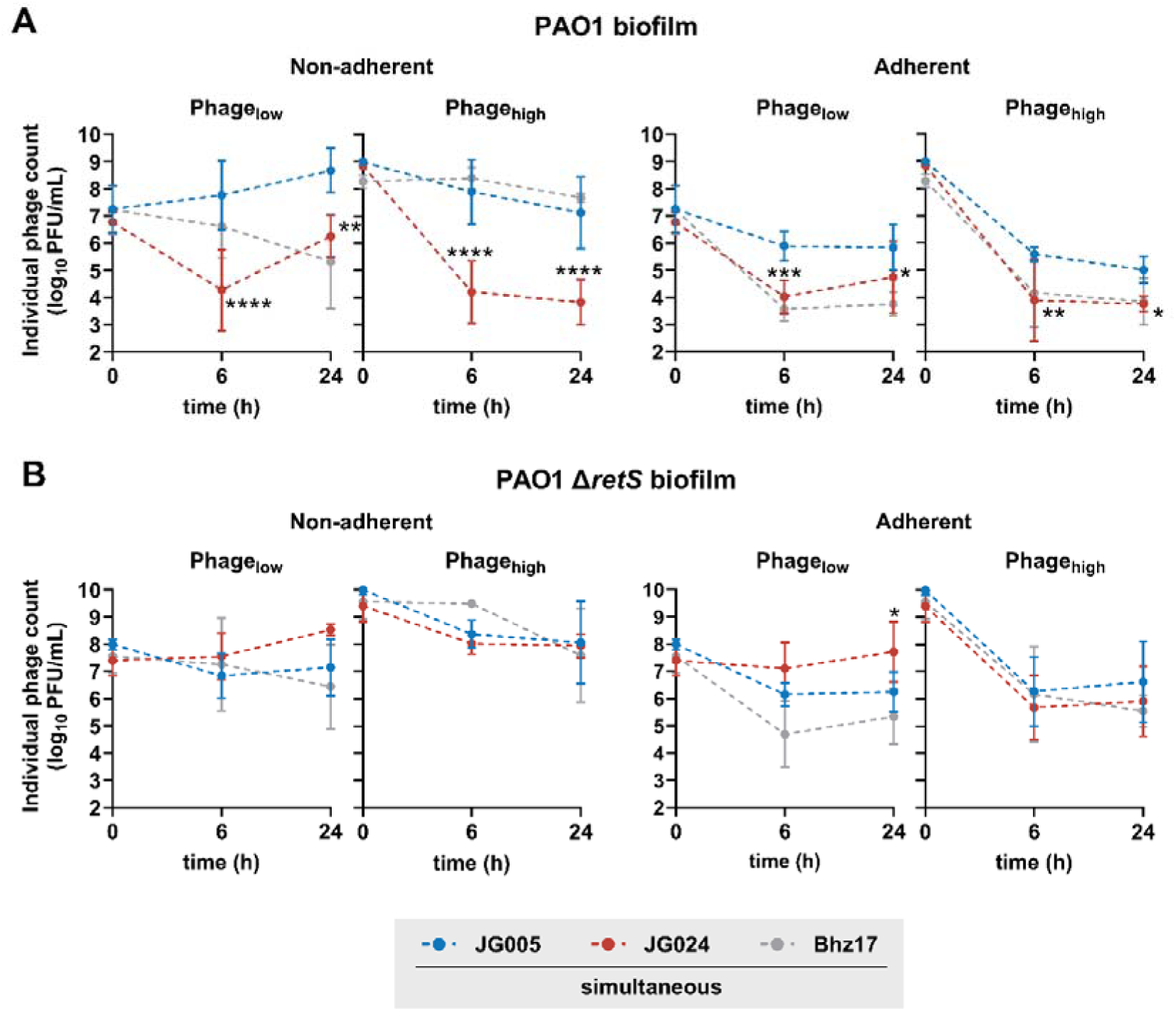
Individual phage replication kinetics of JG005, JG024, and Bhz17 within **(A)** *P. aeruginosa* PAO1 wild-type biofilms and **(B)** PAO1 Δ*retS* biofilms are shown. In the left panels, the number of individual phages of the non-adherent compartment are plotted. Data related to adherent biofilm compartment are shown in right panels. Data represent the mean ± SD of *n*=3 biological replicates with technical duplicates. Individual phage titre from simultaneous treatments were assessed by using specific *P. aeruginosa* indicator strains (**see supplementary methods**). Statistical significance was assessed using Two-way ANOVA and Dunnett’s multiple comparisons test to determine *P*-values, comparing 6- and 24-hour time point (not significant (*P* > 0.05) is not shown, * *P* ≤ 0.05, ** *P* ≤ 0.01, *** *P* ≤ 0.001, **** *P* < 0.0001). Note Bhz17 is unable to replicate in the host PAO1 strain and is therefore serving as a non-replicative control.

Surprisingly, no defect in JG024 fitness was detectable upon examination of phage titre after combined treatment of Δ*retS* biofilm phases: at the phage_high_ concentration its abundance was comparable to that of JG005, while at the phage_low_ concentration it was found to have even increased **(Fig. 3B).** This may explain the more sustained killing at phage_low_ concentrations particularly in the non-adherent phase **(see Fig. 1E-F).** Together, these data show that the interaction of JG005 and JG024 varies corresponding to biofilm phases, bacterial gene function and phage-to-bacteria ratio.

### The latency of JG024 is prolonged in the presence of JG005

To better understand the negative impact of phage JG005 on the replicative fitness of phage JG024, we wondered if these results could be explained by competition for access to the likely shared lipopolysaccharide (LPS) receptor [21–23]. Contrary to this notion, the adsorption of JG024 was not impaired, but rather better than that of JG005, which took an additional 4 minutes to reach 85% threshold when co-administered with JG024 **(Fig. 4A)**.

**Figure 4.**
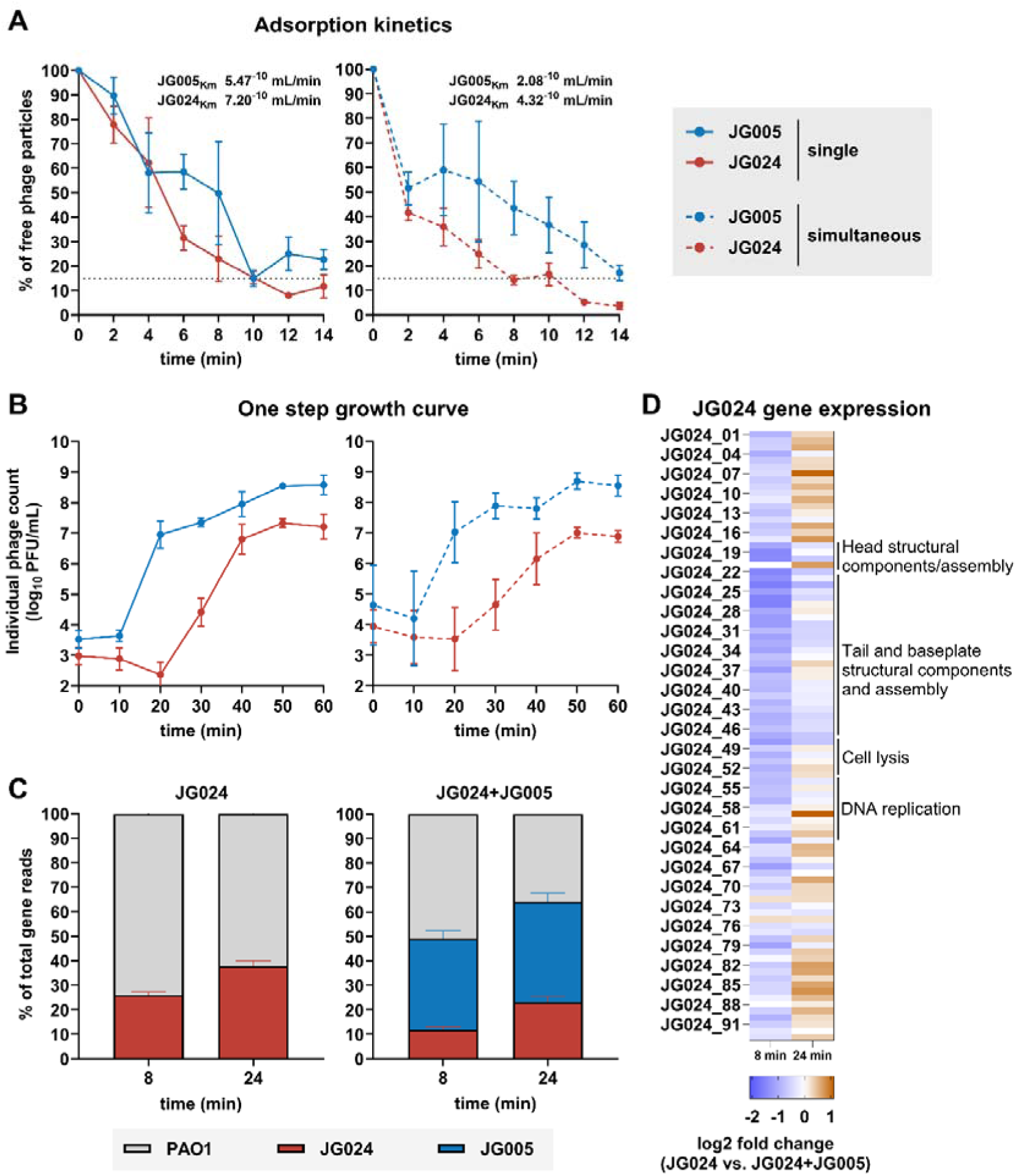
Determination of growth parameters and transcriptional features of phages during coinfection. **(A)** Adsorption kinetics of JG005 and JG024 applied singular or in combination to *P. aeruginosa* PAO1 receptors. Adsorption constant (Km) was calculated from the slope of a linear regression of natural-log free phage particles. The dotted line in the graph indicates the point at which approximately 85% phages were bound (in coinfection: 10 min for JG024, and 14 minutes for JG005). **(B)** One-step growth curve of phages were applied singular or in combination. Solid lines indicate single phage treatment, and dotted lines indicate simultaneous treatment. Data represent the mean ± SD of n=3 biological replicates. **(C)** Percentage of the total gene reads corresponding to PAO1 (grey) and phage gene reads (JG005 – blue, JG024 – red) from dual RNA-seq of PAO1 infected with JG024 or the combination JG024+JG005 at 8 and 24 minutes. RNA-seq data is represent by the mean + SD. **(D)** JG024 transcripts abundance during single or JG024+JG005 coinfection at 8 and 24 minutes after infection. The represented RNA-seq data are generated from two biological replicates.

Both phages are presumed to inject their DNA into PAO1 after adsorption, followed by synthesis of new phage DNA and protein coat, culminating in the lysis of the host. By defining these events by one-step growth curves, we calculated that when added individually, 72 and 4 particles of JG005 and JG024, respectively, were released after 50 minutes of infection **(Fig. 4B)**. During coinfection (with initial input titre of each phage the same as in single infection), the net increase in JG024 progenies was lower, even though the initial infection rate of both phages increased by 10-fold. Underlying reasons for this observed increase in coinfection rate remain unclear. A previous study however showed that coinfection can induce the transcription of gene(s) that encode the cognate bacterial receptor(s), which in turn maximizes the adsorption of temperate phages [38]. Furthermore, JG005 lysis time and number of progenies were largely unaffected by coinfection with JG024 **(Fig. 4B)**. These observations were reproduced with the phylogenetically similar phage pairs, namely JG004 and PTLAW1 (**Fig. S3D-E)**, suggesting that the shorter latency and increased burst size of JG005 allow it to rapidly lyse PAO1.

### JG005 depletes host resources prior JG024 maturation and release

Certain viral species are able to prevent the replication of other viral particles that subsequently infect the same host, a phenomenon referred to as “superinfection exclusion” [39]. We tested this possibility by infecting PAO1 with high titres of JG024 and subsequently with JG005. As shown in **Fig. S4A-B**, JG024-infected PAO1 was found to be permissive for subsequent infection and replication of JG005, albeit the kinetics of both phages appear reduced. In contrast, JG005-infected PAO1 significantly decreased the number of released JG024 particles upon successive infection. Of note, the initial drop in JG024 counts suggests its adsorption and DNA injection processes are unperturbed. There may bespecific JG005-related molecular factor/s disrupting the assembly of JG024 particles. However, based on these data, we rather prefer the notion that the faster replication cycle of JG005 limits the availability of host resources for JG024, which require a longer latency and maturation period, thereby stalling its replication. Furthermore, it is possible that the JG005-JG024 competition may occur not only in the coinfected PAO1 fraction, but also in some of the JG024-only-infected ones, which may then be reinfected and rapidly lysed by JG005.

### Transcriptional response correlates with early dominance of JG005 over JG024

To understand the molecular basis of JG005-JG024 competition, we next compared their genomes. This comparison revealed that JG005 encodes for 88 more genes with at least 15 tRNAs than tRNA-deficient JG024, thereby possibly contributing to shorter latency time of JG005 **(Fig. S5A-B)**. However, despite JG005 encodes additional genetic components its replication, as well as JG024, is impaired in RNA polymerase-inhibited PAO1 (rifampicin treated). No such defects were observed in the solvent control, implying that the respective phages are strictly dependent on host transcriptional components **(Fig. S5C)**.

We subsequently performed dual RNA-Seq to find any potential mechanism by which JG024 assembly is specifically disrupted by comparing the transcript abundances between uninfected, JG024-infected, and JG005+JG024-coinfected PAO1 planktonic culture **(Fig. 4C-D, Dataset S1)**.

As expected, PAO1 transcripts were reduced more under JG024+JG005 coinfection conditions **(Fig. 4C)**. Compared to single infection data, the number of JG024 reads dropped by about 54% and 38% at 8 and 24 minutes, respectively when coinfected with JG005. The observed transcriptome patterns appear to demonstrate a robust correlation with phenotypic data, indicating that JG005 manifests accelerated replication kinetics. When compared with the JG024-alone data, the expression of JG024 genes in JG024+JG005 coinfected samples was reduced at 8 minutes. In contrast, the expression levels of the majority of JG024 genes were found to be comparable between single and coinfected samples at 24 minutes, suggesting a transcriptional recovery **(Fig. 4D)**. When this is correlated with the consistent decline observed for JG024 counts **(see Fig. 2A)**, it indicates that the JG005-mediated bacterial lysis depleted the resources necessary for subsequent steps involving the packaging and release of JG024 virions.

A total of 17 genes were found to be up-regulated between JG024 and JG024+JG005 infected PAO1 compared to uninfected control at both time points. These include *lecA*, phage shock proteins (*PA3728-PA3732*), cyclic diguanylate-regulated protein secretion (*PA4623-PA4624*), bacterioferritin B, and glutaryl-CoA dehydrogenase. Among the 6 down-regulated genes were those of the alkaline protease secretion system (*aprE* and *aprI*), a predicted lipid carrier protein. In addition, particularly at 24 minutes post JG024 or JG024+JG005 infection, transcription of genes associated with energy generation, hydrogen cyanide synthesis and several nutrient transport systems were found to be downregulated, suggesting the onset of phage-mediated metabolic downshift at this time point.

Interestingly, a unique subset of 149 genes were found to be differentially expressed only in JG024-infected PAO1 at 8 or 24 minutes post infection. Among these, an increased expression of the potassium (*kdpA, kdpF, kdpK*), pyoverdine-iron importers (*PA4364-PA4365*), ribonucleotide reductases, fimbriae synthesis, and a decreased expression of outer membrane proteins porin (*oprB*) and of some heme biosynthesis genes. Most of these JG024-specific transcriptional signatures were absent in JG024+JG005 coinfected samples. The negative regulators of alginate synthesis, *wzz* (O-antigen chain length regulator), and *PA909* (initiates autolysis) were among the genes increasingly expressed. Expression of genes involved in nutrient transport and in signaling pathways (e.g., c-di-GMP phosphodiesterase) were found to be decreased at both time points. Taken together, the reduced JG024-specific gene transcription signatures and the increased expression of membrane and metabolic stress-associated genes in PAO1 coinfected with JG024+JG005 strongly suggest that JG005 rapidly exhaust host resources, thereby impairing the ability of JG024 to complete its life cycle **(Dataset S1)**.

### Mutations in a LPS biosynthesis gene confer cross-resistance to JG005 and JG024

PAO1 was found in previous assays to readily evade phage killing (referred here as “bacteriophage insensitive mutants”, short BIMs). We reasoned that the nature and frequency of BIMs could explain differences in the killing capacity of the respective phages and phage-phage competition. Therefore, frequency of BIMs in PAO1 wild–type and PAO1 Δ*retS* was tested in the presence of JG005 and JG024, either alone or in combination. The frequency of JG024 phage-insensitive mutants was 10-fold higher in the Δ*retS* strain than in the wild-type bacteria. However, the BIMs to JG005 or combination treatment (JG005+JG024) arose at a comparable frequency of approximately 10^−7^. Notably, the frequency of JG024-insensitive mutants was higher than that of JG005-treated samples for the Δ*retS* strain. The clinical isolate CHA was tested for comparison and the frequency of mutants was found to be low in response to JG024 or JG024 and JG005 combination **(Figure S6A)**.

We then randomly selected 30 PAO1 BIMs after exposure to each phage, or the combination. Of these, only 18 and 11 variants under the selection pressure of JG005 and JG024, respectively, showed heritable phage resistance. Based on susceptibility profiles, the genomes of 15 representative clones were sequenced for identifying phage resistance mutations, including seven JG005 resistant, six JG024 resistant and two resistant to both phages **(Fig. S6B, Table S1)**.

Four sequenced JG005 resistant PAO1 mutants had mutations in the *wzy* gene, which encodes B and O antigen polymerase required for LPS synthesis. The remaining two had missense mutations in *algC* (essential for LPS processing) or *lolC* (encodes lipoprotein localization protein) and were found to reduce susceptibility to JG005. In contrast, among the six mutants resistant to JG024, three had a missense or frameshift mutation in *wzy,* while the remaining three had a missense or nonsense mutation in *wzz2*, a gene that regulates O-antigen chain length. Finally, both mutants selected on medium containing the JG005+JG024 combination solely contained a frameshift mutation in *wzy*. This analysis indicates that a mutation in *wzy* confers cross-resistance to both JG005 and JG024 and that the degree of selective pressure exerted by each phage is different.

Moreover, to test whether the phage-resistance conferring mutation is strain dependent, we sequenced the genome of three JG024-resistant clones from PA14 strain background. Notably, these mutants had a missense mutation in the *ssg* gene, which is also involved in LPS synthesis. This observation suggests that the genetic determinants involved in JG024 resistance may vary depending on the genetic background of *P. aeruginosa*.

### Sequential administration prevents JG005-JG024 competition

In planktonic conditions over 24 hours, when JG024 was added 2 hours before JG005, bacterial growth tended to be slightly more inhibited **(Fig. 5A)**, and again, without affecting JG024 replication **(Fig. 5B)**. While the reduction in bacterial viability was comparable between sequential **(Fig. 5C-D)** and simultaneous phage treatment **(see Fig. 1C)**, the phage-phage competition in both biofilm phases was circumvented with sequential phage treatment **(Fig. 5E-F)**.

**Figure 5.**
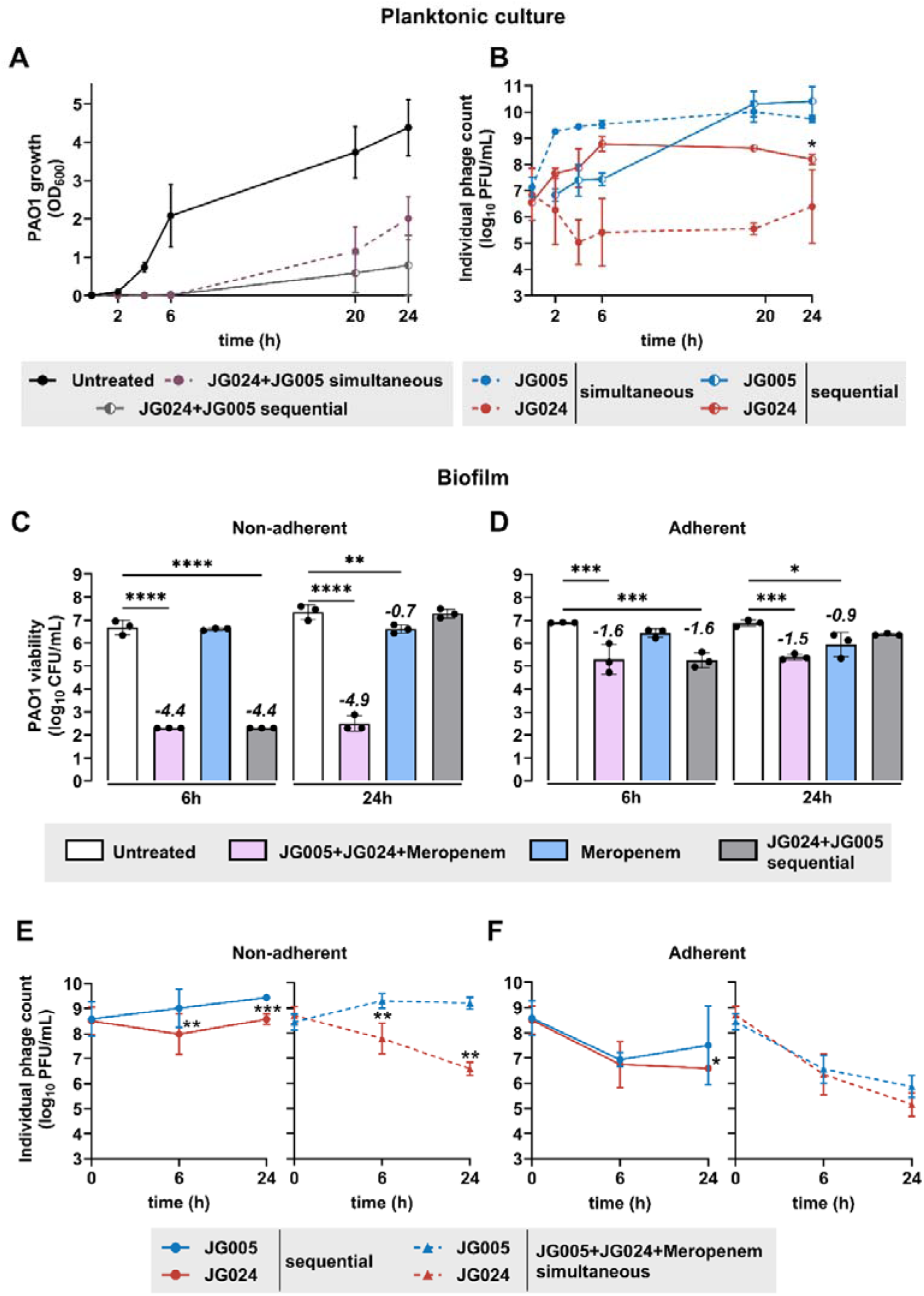
Sequential phage administration and phage and antibiotic combinations effects against *P. aeruginosa* biofilm. For simultaneous and sequential phage application in planktonic culture both phages were added either simultaneously or JG024 was applied sequentially 2 hours before JG005. **(A)** Bacterial growth in response to simultaneous (purple) or sequential (grey) phage treatment of *P. aeruginosa* PAO1 planktonic culture. **(B)** Phage replication patterns of JG024 (red) and JG005 (blue) when added to PAO1 planktonic culture. Simultaneous treatment is in dotted lines, and sequential phage administration is in solid lines. Statistical significance was determined using One-way ANOVA and Šídák’s multiple comparisons test, comparing simultaneous and sequential treatment conditions at 24-hour time point, * *P* < 0.05. Bacteria and phage counts of PAO1 biofilms treated either sequentially with phages, or both phages concomitantly with meropenem were enumerated at 6- and 24-hours timepoint. Bacterial viability of non-adherent population **(C)** and adherent **(D)** compartment of static PAO1 biofilm of sequential phage treated (first JG024, two hours later JG005 in grey), simultaneous phage combination together with meropenem (5 µg/mL, in light pink), and meropenem alone (5 µg/mL, in light blue). The corresponding phage viability of both non-adherent **(E)** and adherent **(F)** compartments is plotted. Sequentially administered phages are shown in solid lines (JG005 blue, JG024 red) and a simultaneous combination with meropenem in dotted lines and triangles. Data represent the mean ± SD of *n*=3. Numeric values indicate log-fold changes relative to the untreated control. Statistical significance was assessed using Ordinary One-way ANOVA to determine *P*-values for bacterial viability and Two-way ANOVA and Šídák’s multiple comparisons test comparing 6 and 24-hour time point for phage viability (not significant (*P* > 0.05) is not shown, * *P* < 0.05, ** *P* < 0.01, *** *P* < 0.001, **** *P* < 0.0001). Detection limit is 200 CFU/mL, all samples below are set to this limit.

### Concomitant treatment with phages and meropenem restricts biofilm dispersion

The synergistic effects of phage and antibiotics have been previously documented [5–7, 40]. To test the impact of 5X minimum inhibitory concentration of meropenem (frequently used for treating severe infections in critically ill patients) on phage-mediated killing and JG005-JG024 competition in biofilm, we included two treatment groups: phages and meropenem added simultaneously, as well as meropenem alone. No reduction of bacterial growth was observed in the meropenem-only group. However, PAO1 biofilm dispersal was restricted over 24 hours when meropenem and phage combinations were added simultaneously **(Fig. 5C)**. This effect was only detected in the non-adherent phase, while only a moderate reduction of bacterial viability at 24 hours was observed in the adherent phase **(Fig. 5D)**. The addition of meropenem however did not alleviate JG024-JG005 competition, particularly in the non-adherent phase where JG024 was found to be reduced **(Fig. 5E-F)**.

## Discussion

In this study, we incorporated additional readouts to conventional microtitre plate-based biofilm assays, demonstrating that the efficacy of phage combination varied in correspondence with distinct phases of *P. aeruginosa* biofilm: while surface-attached biofilm exhibited tolerance, the dispersed biofilm population was relatively susceptible, yet subsequently allowed for the emergence of phage resistance. Such spatiotemporal differences in phage effects can be observed not only in biofilms but also in soil, the plant rhizosphere and the gut lumen [41–43]. Diffusion-limiting structures tend to restrict the free interaction between phage(s) and bacteria and/or promote their coexistence [44–47]. Additional factors which might contribute to the recalcitrance of *P. aeruginosa* PAO1 biofilms to phage lysis have been previously described: a subpopulation of bacteria in biofilms tend to lose B-band and O-antigen of LPS, which is the putative receptor for JG005 and JG024. Slow-growing bacterial cells then undergo metabolic changes including cyclic AMP production, reducing their susceptibility to phage infection and lysis [48, 49]. Finally, transcriptional counter-responses or mutation acquisition under phage-induced stress can influence subset of genes or select hypermorphic mutations that increase biofilm formation, thereby overturning the initial phage-mediated anti-biofilm effects [50, 51]. Nevertheless, our finding that meropenem and JG005/JG024 combination treatment strongly inhibited the dispersal of resistant/tolerant biofilm population to both agents is encouraging. This also mirrors the findings of some clinical case reports that showed better treatment efficacy when phage and antibiotic combinations were used [52]. Therefore, the optimized *in vitro* assay readouts here would allow reliable selection of therapeutic phages and/or antibiotic combinations that can most effectively target more than one metabolic state of PAO1.

Our assay design enabled us to detect phage-phage competition. Although the titres of JG005 and JG024 were matched prior to simultaneous treatment, their ability to coinfect, assemble structures and complete their lytic cycle was found to be asymmetric. As a result, replication of the “weaker” phage was inhibited, especially at phage_high_ ratios. This observation is reminiscent of “interspecific competition”, a phenomenon that occurs across different species, including eukaryotic viruses coinfecting plants or animals [53]. While there are different contexts and conditions that can lead to competition, intrinsic features of the phage itself are a major determinant [54–56]. For example, dsDNA phages, with a short latency period and a high burst size, encoding their own set of tRNAs may reduce the dependence of phages on the bacterial host to synthesize their macromolecules as well as allowing evasion of tRNA-targeting host defenses. Fulfilling these criteria, the JG005 genome harbors twice as many genes as JG024, including 15 tRNAs, which likely contribute to its superior replicative fitness and its respective dominance over JG024. The resulting asymmetric selection pressure exerted by each phage appeared to alter the spectrum of phage resistance mutations in our study, as reported previously for other phages and clinically approved drugs [57, 58]. At early time points in our experiments, the selective pressure on PAO1 was likely to have been exerted predominantly by JG005, leading to resources monopolization, and strong selection for JG005 mutants, which also showed cross-resistance to JG024, leading to further attenuation of JG024 replication.

Nevertheless, our observations in the *P. aeruginosa* strain CHA, where the initial impairment in JG024 replication was subsequently restored, cannot be solely explained by this model. We have shown that the frequency of phage-insensitive mutants was strikingly lower in the clinical CHA strain than in PAO1 strain. This may have reduced the conflicts in replication of JG005 and JG024 and/or with the different underlying mechanisms of phage resistance in CHA strain. Reinforcing this view, our analysis showed that gene mutation conferring JG024 phage resistance is not identical in *P. aeruginosa* strains PAO1 and PA14. Such strain-specific irregularities in the mutational spectrum are not uncommon [59]. Thus, *P. aeruginosa* genotype (and specific anti-phage defense systems) may influence the strength of selective pressure exerted by individual phages, thereby altering dynamics of bacterial adaptation, phage resistance evolution and phage-phage competition.

Sequential administration of JG024 and JG005 reduced phage-phage competition but did not achieve sustained bacterial killing, nor prevented phage resistance in biofilms. Wright *et al.* [57] showed that only sequential addition of two LPS-binding phages resulted in the acquisition of several resistance mutations in *P. aeruginosa*. Also, Hall *et al.* found that the simultaneous application of four phages was either equally effective or even superior to sequential application in reducing *P. aeruginosa* viability and in mitigating phage resistance [60]. As we used different phages and bacterial strains, direct comparison of our results with others is not appropriate. Nevertheless, phages targeting different bacterial receptors with similar replication features may constitute a compatible phage combination minimizing cross-resistance and phage-phage competition.

Inherent limitations of our experiments are the short period of observation and the inability to evaluate other confounders like immune response, phage administration route and schedule. Our results thus probably overestimate the frequency and significance of phage resistance and phage-phage competition compared to the context of human disease. The range, strength and course of phage-bacteria-host immune interactions are expected to differ among patients, and even within the same patient, collectively influencing the outcome. For example, the phage resistant mutants deficient in the production of the key virulence factor LPS rapidly evolve under laboratory conditions, but may not emerge/survive long in immunocompetent patients receiving phage therapy [52]. Future investigations should encompass heightened complexity recapitulating more clinically pertinent conditions in evaluating phage combinations. In this regard, animal models are useful for establishing the safety and systemic availability of phage therapeutics and for understanding the implications of metazoan immunity on phage-bacteria interactions [41, 61–64]. However, their routine use in identifying the most effective phage combinations against each patient isolate is limited. Therefore, it is imperative to employ a suite of *in vitro/ex vivo* assays with expanded readouts to evaluate key variables pertaining to human immunity, pathogen, and the specific phage. Data from such laboratory assays may provide greater assurance that a selected phage combination is optimal for clinical use.

## Author Contributions

**Conception and design: M.B., G.N., J.-M.G., A.C.H., K.P., L.D., J.-D.R., M.W., S.-M.W, G.K.**

**Funding: G.N., J.-M.G., A.C.H., K.P., L.D., J.-D.R., M.W., S.-M.W., G.K.**

**Data Acquisition, formal analysis: M.B., I.H.E.K., A.L., S.K., C.W., X.M.T., B.G., G.K.**

**Data interpretation: M.B., I.H.E.K., G.N., J.-M.G., A.C.H., K.P., L.D., M.W., S-M.W., G.K.**

**Writing – original draft: M.B., G.K.**

**Writing – revision and editing: all authors**

## Acknowledgments

This work received funding from the Agence Nationale de la Recherche, France and Bundesministerium für Bildung und Forschung, Germany under the collaborative grant “MAPVAP”, FKZ 01KI2124 (to G.N., A.C.H., M.W., S.-M.W. and G.K.) and ANR-19-AMRB-0002 (to J.-D.R., L.D. and J.-M.G.) and from Mukoviszidose Institut GmbH, Bonn, the research and development arm of the German Cystic Fibrosis Association Mukoviszidose e.V, project number P2201 (to M.W., K.P., G.N., and G.K.). K.P. acknowledges support from the DFG (SPP2389, project-ID 503931087 and EXC 2051, project-ID 390713860) and the European Research Council (ArtRNA, CoG-101088027).

Authors thank Monot Marc, Biomics Platform, C2RT, Institut Pasteur, Paris, France, supported by France Génomique (ANR-10-INBS-09) and IBISA for sequencing phage-resistant clones, and Maren Mieth for A549 cells, the Fraunhofer Institute for Toxicology and Experimental Medicine, especially Sarah Wienecke and Holger Ziehr, for purified phage preparation, Johannes Wittmann for JG005 phage genome, Susanne Häußler for *P. aeruginosa* strains CH3549 and F2230, Matthew Parsek and Alain Filloux for *P. aeruginosa* strains PAO1 Δ*retS* and Rike Zietlow for editorial help.

---

**Figure S1.**
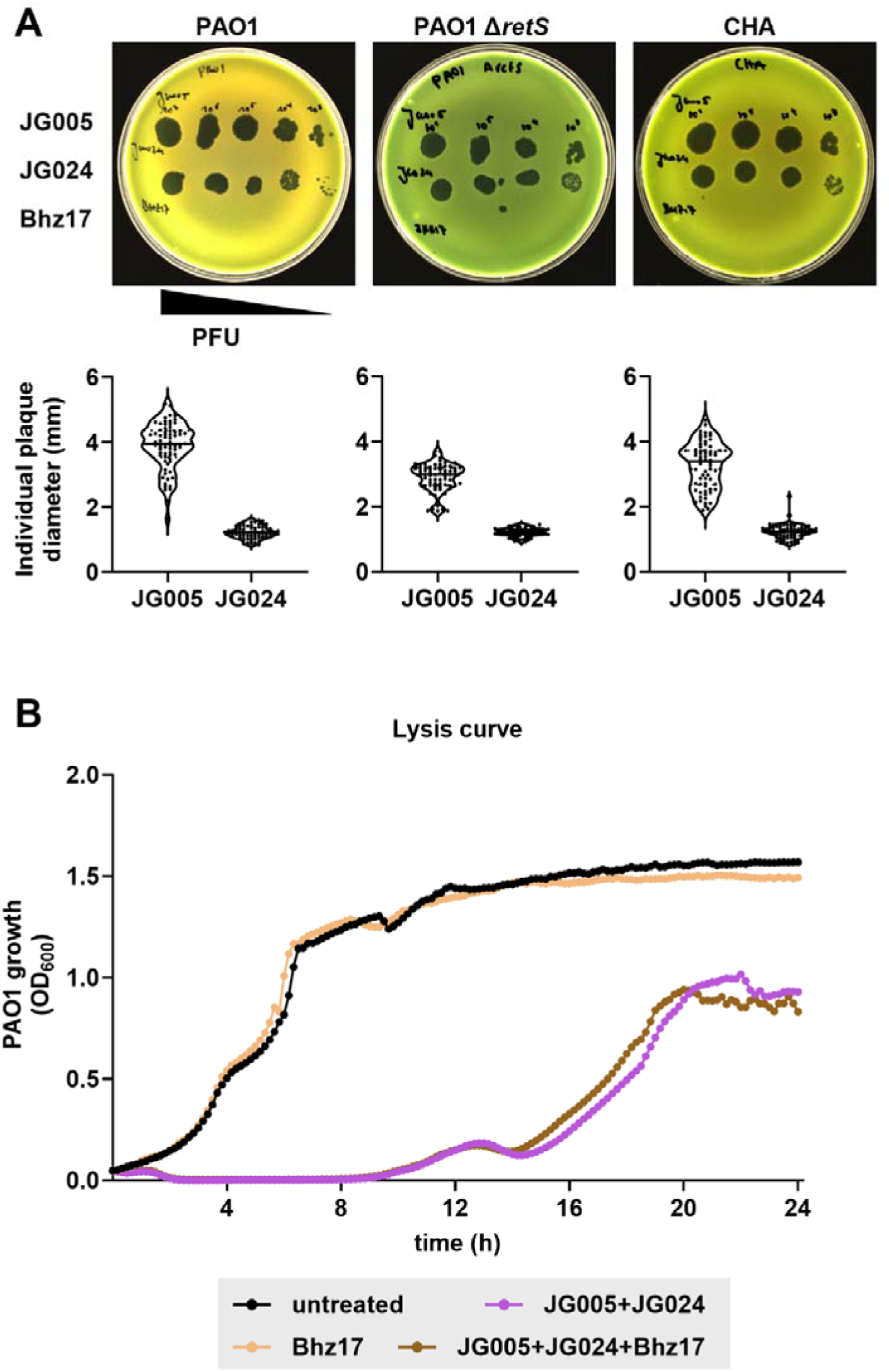
Phages and their host specificity. **(A)** Strain-specific variation in *P. aeruginosa* phage susceptibility. Phage host range was assessed by adjusting phage lysates to 1E6 PFU/mL and serially diluting them 10-fold. The dilutions were spotted onto agar overlays (4 μL per spot) prepared with the indicated *P. aeruginosa* strains (PAO1, PAO1 Δ*retS*, CHA). At least 70 plaques for each *P. aeruginosa* strain were selected for measuring the individual size using the plaque size tool (55). Data are represented as violin blots. **(B)** Representative phage lysis curves of the non-replicative control phage Bhz17 individually (single) and simultaneous combinations of phages JG005 and JG024 with and without Bhz17, on PAO1 at a phage-bacterium ratio of 1:1. Bacterial growth (OD_600_) was measured in plate reader for 24 hours at 37 °C (SpectraMax, *n*=3).

**Figure S2.**
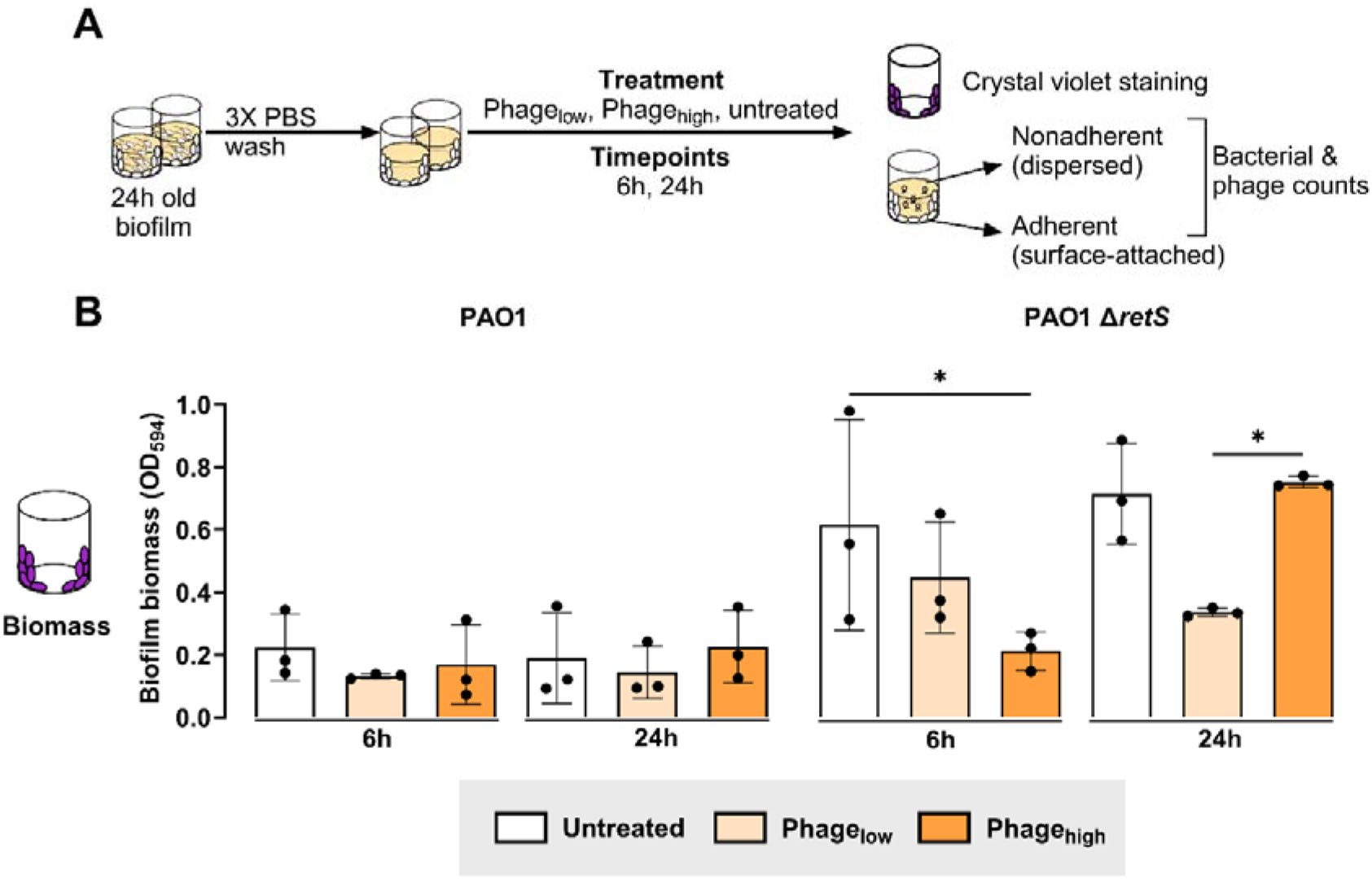
Static *in vitro* biofilm production of *P. aeruginosa* PAO1 wild-type and PAO1 Δ*retS*. **(A)** Schematic illustration of biofilm assay: An equimolar mixture of three *P. aeruginosa*-specific phages was tested in a 10-fold low/high phage:bacteria ratio, on 24 hours-old *P. aeruginosa* PAO1 static biofilms in microtitre plates. Separated adherent (surface-attached) or non-adherent (dispersed) biofilm phases were used to determine bacterial and phage counts. Biomass of adherent biofilm was stained with crystal violet and measured at OD 590 nm. **(B)** Crystal violet staining of biofilm biomass of the adherent compartment of PAO1 wild-type and PAO1 Δ*retS*. Data represent the mean ± SD of *n*=3 biological replicates from technical triplets. Statistical significance was assessed using Ordinary One-way ANOVA and Šídák’s multiple comparisons test to determine *P*-values (not significant (*P* > 0.05) is not shown, * *P* < 0.05).

**Figure S3.**
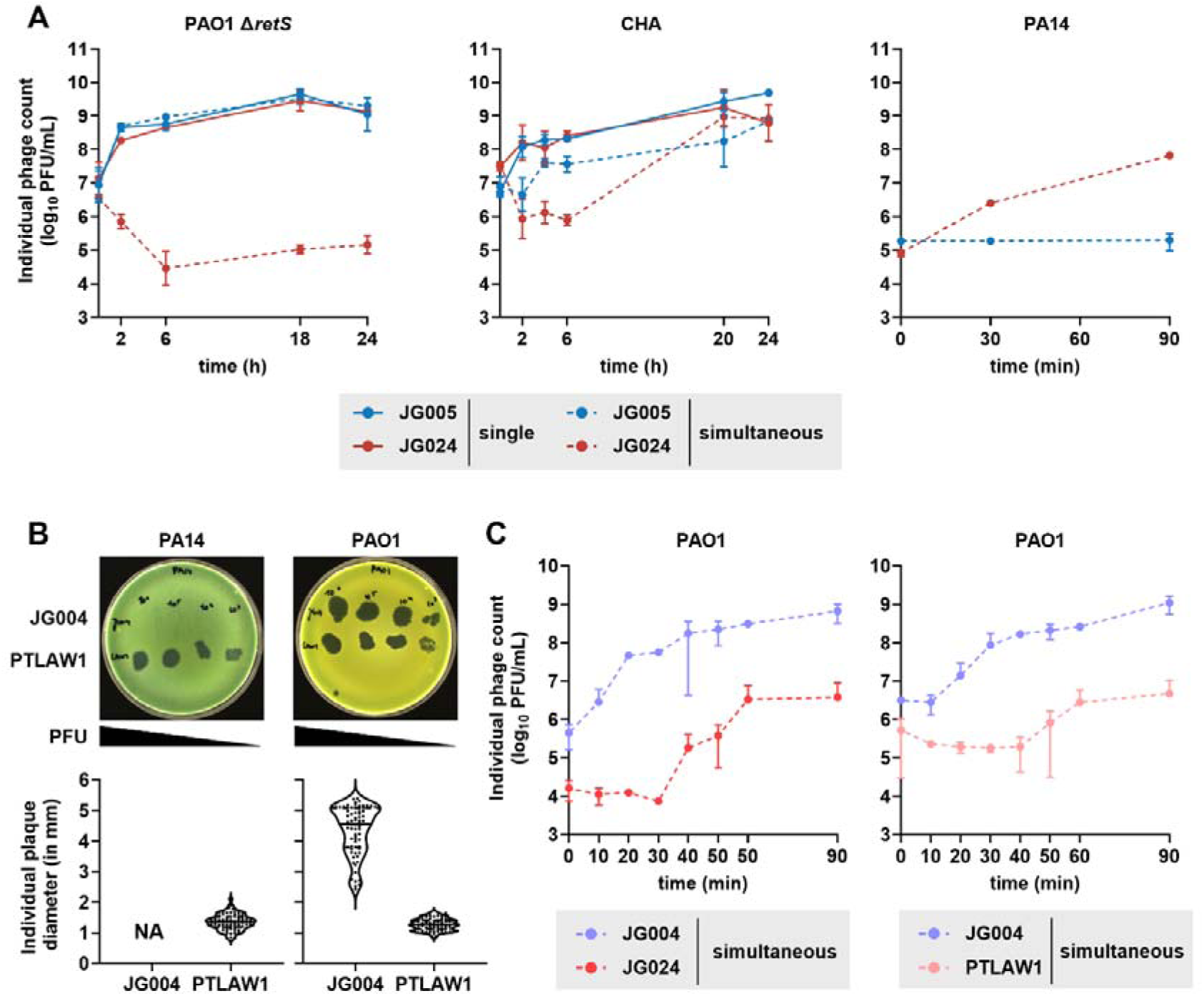
Replication and growth parameters of each phage in different *P. aeruginosa* strains during single infection or coinfection. **(A)** Individual phage counts in PAO1 Δ*retS*, clinical isolate *P. aeruginosa* CHA, and in PA14 strain. Note JG005 cannot replicate in the host PA14 strain. **(B)** Phages JG004 and PTLAW1 are phylogenetically similar to JG005 and JG024, respectively. Both show comparable host range and plaque size on the indicated strains. Plaque size was calculated from 70 plaques per phage using the plaque size tool (55). Plaque size data are represented as violin plots. **(C)** When simultaneously added with JG004, the kinetics of JG024 and PTLAW1 are delayed. Data represent the mean ± SD of *n*=2 biological replicates.

**Figure S4.**
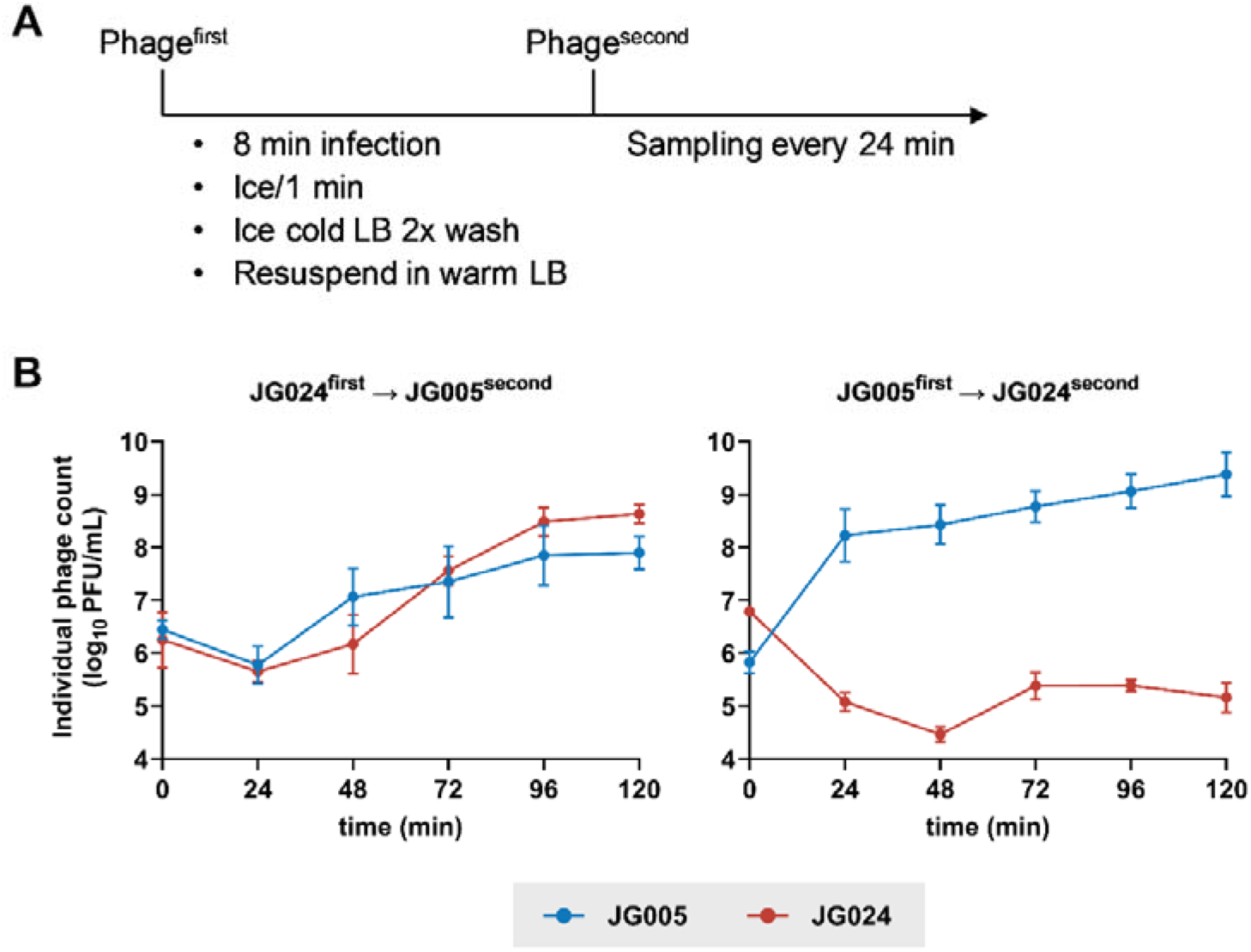
Superinfection exclusion. **(A)** Schematic illustration of the experimental setting, to test if primary phage infection with one phage excludes superinfection with the second phage. **(B)** A pre-established infection of PAO1 with JG005 or JG024at a phage-bacterium-ratio of 10:1 for 8 minutes was superinfected with the respective other phage at a phage-bacterium-ratio of 1:10. Subsequently individual phage production was monitored every 24 minutes for 2 hours. Data represent the mean ± SD of *n*=3 biological replicates.

**Figure S5.**
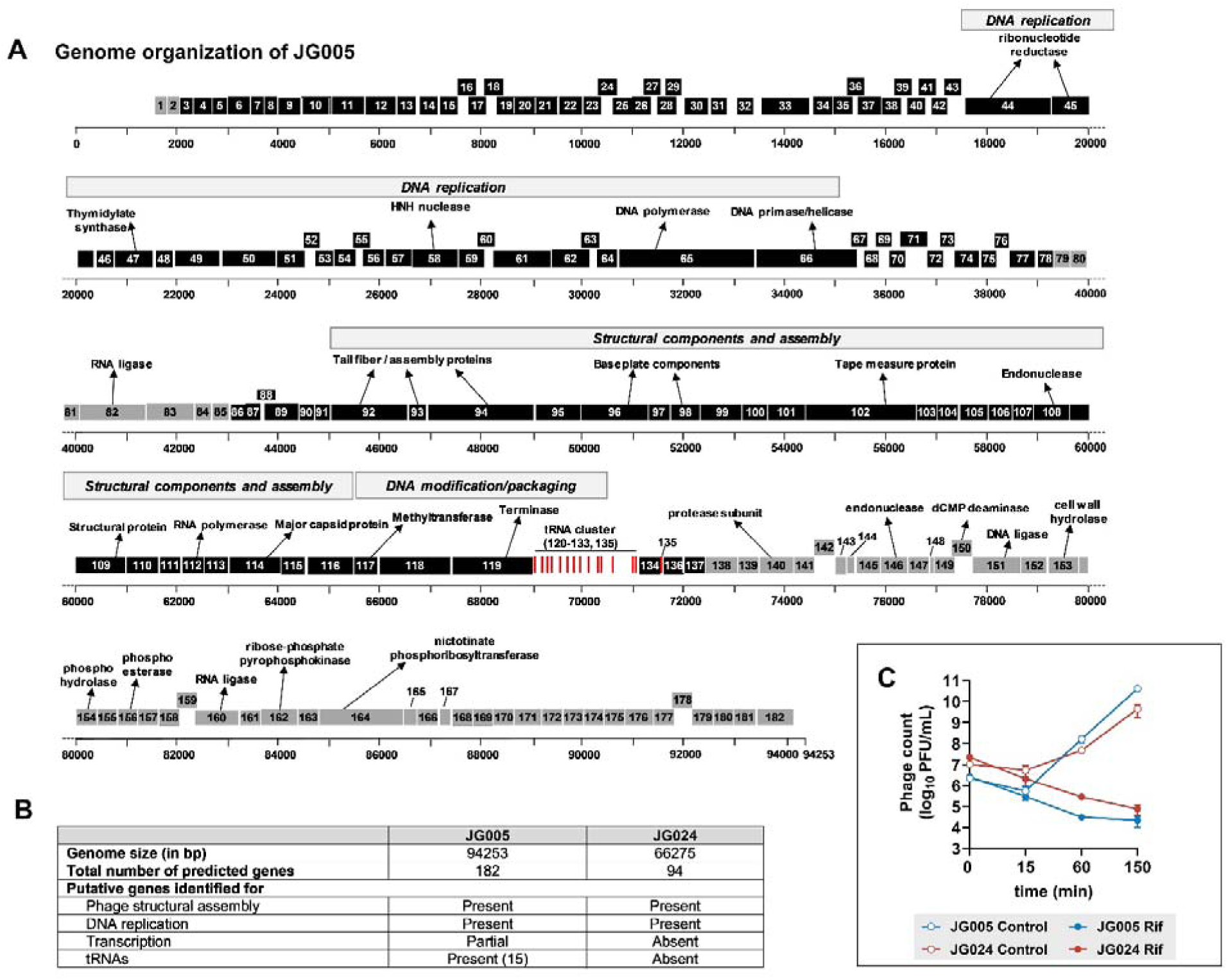
Genomic comparison of JG005 and JG024. **(A)** Predicted coding sequences, the tRNA locations, some functional assignments and overall genome organization of phage JG005. The double-stranded DNA genome of phage JG005 is represented as a horizontal bar with vertical markers at every kilobase pair. Gene numbers are shown in boxes, which are shaded in black or grey according to their transcription from plus and negative strands, respectively. Putative gene functions, based on BLAST analyses, are indicated. A Megablast analysis revealed that JG005 has close similarity to other bacteriophages (JG004, PAK_P1, PAK_P2, PAK_P4, PaP1, and vB_PaeM_C2-10_Ab1) infecting also *P. aeruginosa* PAO1. In particular, phages JG004 and JG005 are almost identical and the difference between their DNA sequences is limited to a few nucleotides. **(B)** Genomic characteristics of JG005 and JG024. **(C)** Both JG005 and JG024 require *P. aeruginosa* transcriptional neries for their replication. Rifampicin-treated PAO1 (Rif) restricts the replication of both phages that are dependent on bacterial RNA erase. Data represent the mean ± SD of *n*=2 biological replicates.

**Figure S6.**
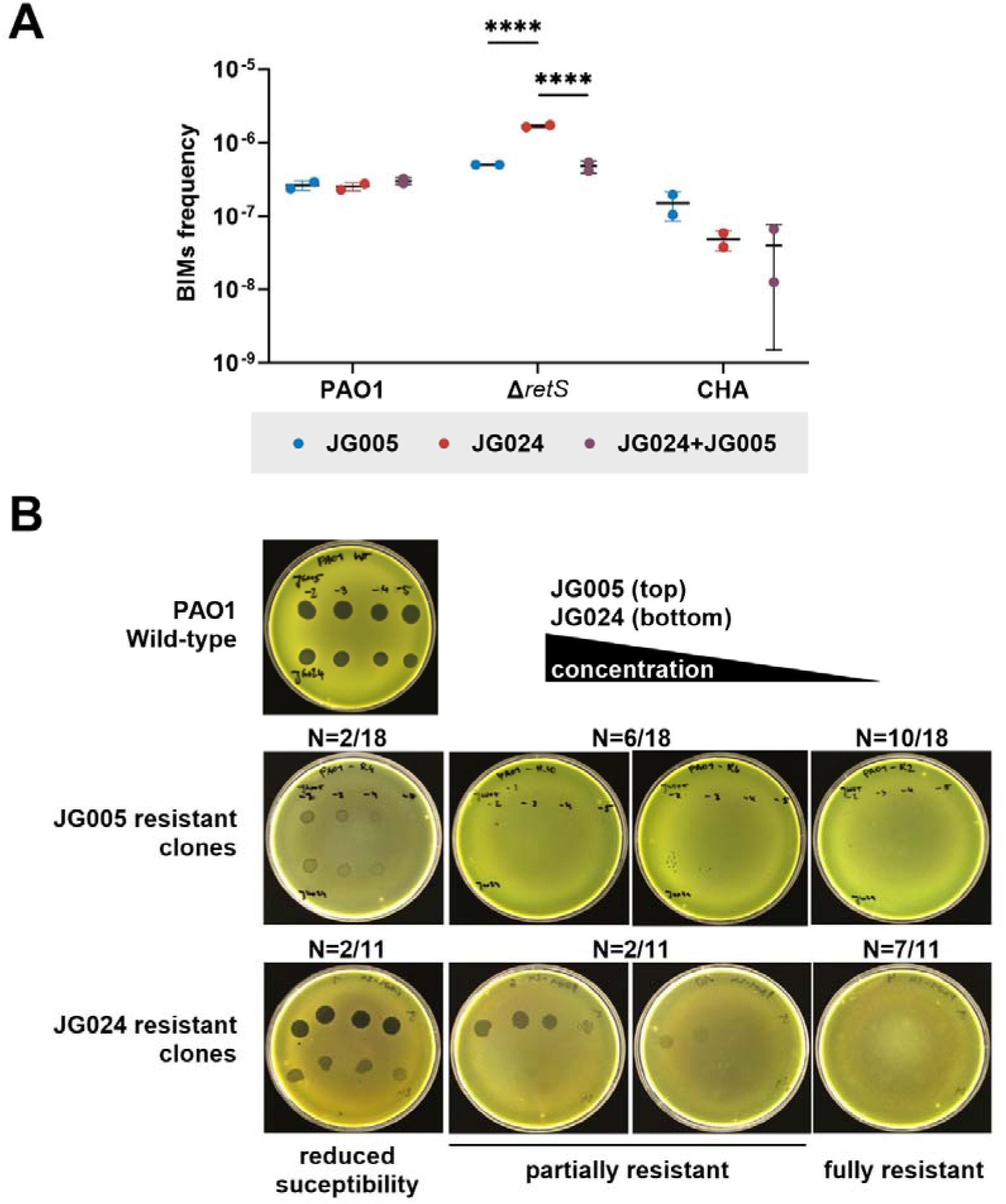
Phage resistance frequency and phenotypic assessment. Frequency of bacteriophage-insensitive mutants (BIM) from *P. aeruginosa* PAO1 wild-type, PAO1 Δ*retS*, and CHA after overnight incubation with the respective phages (phage-to-bacteria ratio 50:1) **(A)** Statistical significance was assessed using Ordinary Two-way ANOVA and Šídák’s multiple comparisons test to determine *P*-values (not significant (*P* > 0.05) is not shown, **** *P* < 0.0001)**. (B)** Representative JG005- and JG024-resistant PAO1 wild-type BIM with altered susceptibility patterns, compared to the indicated wild-type **(see Table S1)**. 10-fold dilutions of both phages were individually spotted on the lawn of the indicated BIM, with JG005 (top) and JG024 (bottom). The phage concentration decreases from left to right, except for the JG024-resistant clones, where the phage concentration decreases from right to left. Therefore, these images were mirrored to match the concentration gradient.

## References

1. Malhotra S, Hayes Jr D, Wozniak DJ. Cystic fibrosis and *pseudomonas aeruginosa*: The host-microbe interface. Clin Microbiol Rev. 2019;32:e00138–18 10.1128/cmr.00138-18

2. Sauer K, Stoodley P, Goeres DM et al. The biofilm life cycle: Expanding the conceptual model of biofilm formation. Nat Rev Microbiol. 2022;20:608–20 10.1038/s41579-022-00767-0

3. Lebeaux D, Ghigo J-M, Beloin C. Biofilm-related infections: Bridging the gap between clinical management and fundamental aspects of recalcitrance toward antibiotics. Microbiol Mol Biol Rev. 2014;78:510–43 10.1128/MMBR.00013-14

4. Marongiu L, Burkard M, Lauer UM et al. Reassessment of historical clinical trials supports the effectiveness of phage therapy. Clin Microbiol Rev. 2022:e00062–22 10.1128/cmr.00062-22

5. Segall AM, Roach DR, Strathdee SA. Stronger together? Perspectives on phage-antibiotic synergy in clinical applications of phage therapy. Curr Opin Microbiol. 2019;51:46–50 10.1016/j.mib.2019.03.005

6. De Soir S, Parée H, Kamarudin NHN et al. Exploiting phage-antibiotic synergies to disrupt *pseudomonas aeruginosa* pao1 biofilms in the context of orthopedic infections. Microbiol spectr. 2024;12:e03219–23 10.1128/spectrum.03219-23

7. Comeau AM, Tétart F, Trojet SN et al. Phage-antibiotic synergy (pas): Β-lactam and quinolone antibiotics stimulate virulent phage growth. PLoS One. 2007;2:e799 10.1371/journal.pone.0000799

8. Roach DR, Leung CY, Henry M et al. Synergy between the host immune system and bacteriophage is essential for successful phage therapy against an acute respiratory pathogen. Cell Host Microbe. 2017;22:38–47. e4 10.1016/j.chom.2017.06.018

9. Georjon H, Bernheim A. The highly diverse antiphage defence systems of bacteria. Nat Rev Microbiol. 2023;21:686–700 10.1038/s41579-023-00934-x

10. Palmer GC, Whiteley M. Metabolism and pathogenicity of *pseudomonas aeruginosa* infections in the lungs of individuals with cystic fibrosis. Microbiol Spectr. 2015;3:185–213 10.1128/microbiolspec.MBP-0003-2014

11. Golden MM, Post SJ, Rivera R et al. Investigating the role of metabolism for antibiotic combination therapies in *pseudomonas aeruginosa*. ACS Infect Dis. 2023;9:2386–93 10.1021/acsinfecdis.3c00452

12. Manner C, Dias Teixeira R, Saha D et al. A genetic switch controls *pseudomonas aeruginosa* surface colonization. Nat Microbiol. 2023;8:1520–33 10.1038/s41564-023-01403-0

13. Meneses L, Brandão AC, Coenye T et al. A systematic review of the use of bacteriophages for *in vitro* biofilm control. Eur J Clin Microbiol Infect Dis. 2023;42:919–28 10.1007/s10096-023-04638-1

14. Pires DP, Melo LD, Azeredo J. Understanding the complex phage-host interactions in biofilm communities. Annu Rev Virol. 2021;8:73–94 10.1146/annurev-virology-091919-074222

15. Molina F, Menor-Flores M, Fernández L et al. Systematic analysis of putative phage-phage interactions on minimum-sized phage cocktails. Sci Rep. 2022;12:2458 10.1038/s41598-022-06422-1

16. Refardt D. Within-host competition determines reproductive success of temperate bacteriophages. ISME J. 2011;5:1451–60 10.1038/ismej.2011.30

17. Turner PE. Parasitism between coLinfecting bacteriophages. Adv Ecol Res. 2005;37:309–32 10.1016/S0065-2504(04)37010-8

18. Visnapuu A, Van der Gucht M, Wagemans J et al. Deconstructing the phage–bacterial biofilm interaction as a basis to establish new antibiofilm strategies. Viruses. 2022;14:1057 10.3390/v14051057

19. Sutherland IW, Hughes KA, Skillman LC et al. The interaction of phage and biofilms. FEMS Microbiol Lett. 2004;232:1–6 10.1016/S0378-1097(04)00041-2

20. Jault P, Leclerc T, Jennes S et al. Efficacy and tolerability of a cocktail of bacteriophages to treat burn wounds infected by *pseudomonas aeruginosa* (phagoburn): A randomised, controlled, double-blind phase 1/2 trial. Lancet Infect Dis. 2019;19:35–45 10.1016/S1473-3099(18)30482-1

21. Garbe J, Bunk B, Rohde M et al. Sequencing and characterization of *pseudomonas aeruginosa* phage jg004. BMC Microbiol. 2011;11:1–12 10.1186/1471-2180-11-102

22. Garbe J, Wesche A, Bunk B et al. Characterization of jg024, a *pseudomonas aeruginosa* pb1-like broad host range phage under simulated infection conditions. BMC Microbiol. 2010;10:1–10 10.1186/1471-2180-10-301

23. Selezska K, Kazmierczak M, Müsken M et al. *Pseudomonas aeruginosa* population structure revisited under environmental focus: Impact of water quality and phage pressure. Environ Microbiol. 2012;14:1952–67 10.1111/j.1462-2920.2012.02719.x

24. Henry M, Bobay LM, Chevallereau A et al. The search for therapeutic bacteriophages uncovers one new subfamily and two new genera of *pseudomonas*-infecting *myoviridae*. PLoS One. 2015;10:e0117163 10.1371/journal.pone.0117163

25. Trofimova E, Jaschke PR. Plaque size tool: An automated plaque analysis tool for simplifying and standardising bacteriophage plaque morphology measurements. Virology. 2021;561:1–5 10.1016/j.virol.2021.05.011

26. O’flynn G, Ross R, Fitzgerald G et al. Evaluation of a cocktail of three bacteriophages for biocontrol of *escherichia coli* o157: H7. Appl Environ Microbiol 2004;70:3417–24 10.1128/AEM.70.6.3417-3424.2004

27. Chevallereau A, Blasdel BG, De Smet J et al. Next-generation “-omics” approaches reveal a massive alteration of host rna metabolism during bacteriophage infection of *pseudomonas aeruginosa*. PLoS Genet. 2016;12:e1006134 10.1371/journal.pgen.1006134

28. Hyman P, Abedon ST. Practical methods for determining phage growth parameters. Methods Mol Biol. 2009;501:175–202 10.1007/978-1-60327-164-6_18

29. Ramsey J, Rasche H, Maughmer C et al. Galaxy and apollo as a biologist-friendly interface for high-quality cooperative phage genome annotation. PLoS Comput Biol. 2020;16:e1008214 10.1371/journal.pcbi.1008214

30. Okonechnikov K, Golosova O, Fursov M et al. Unipro ugene: A unified bioinformatics toolkit. Bioinformatics. 2012;28:1166–67 10.1093/bioinformatics/bts091

31. Laslett D, Canback B. Aragorn, a program to detect trna genes and tmrna genes in nucleotide sequences. Nucleic Acids Res. 2004;32:11–16 10.1093/nar/gkh152

32. Lowe TM, Eddy SR. Trnascan-se: A program for improved detection of transfer rna genes in genomic sequence. Nucleic Acids Res. 1997;25:955–64 10.1093/nar/25.5.955

33. Ceyssens P-J, Minakhin L, Van den Bossche A et al. Development of giant bacteriophage Lkz is independent of the host transcription apparatus. J Virol. 2014;88:10501–10 10.1128/jvi.01347-14

34. Culviner PH, Guegler CK, Laub MT. A simple, cost-effective, and robust method for rrna depletion in rna-sequencing studies. mBio. 2020;11:e00010–20 10.1128/mbio.00010-20

35. Moscoso JA, Mikkelsen H, Heeb S et al. The *pseudomonas aeruginosa* sensor rets switches type iii and type vi secretion via cLdiLgmp signalling. Environ Microbiol. 2011;13:3128–38 10.1111/j.1462-2920.2011.02595.x

36. Goodman AL, Kulasekara B, Rietsch A et al. A signaling network reciprocally regulates genes associated with acute infection and chronic persistence in *pseudomonas aeruginosa*. Dev Cell. 2004;7:745–54 10.1016/j.devcel.2004.08.020

37. Dacheux D, Toussaint B, Richard M et al. *Pseudomonas aeruginosa* cystic fibrosis isolates induce rapid, type iii secretion-dependent, but exou-independent, oncosis of macrophages and polymorphonuclear neutrophils. Infect Immun. 2000;68:2916–24 10.1128/iai.68.5.2916-2924.2000

38. Joseph SB, Hanley KA, Chao L et al. Coinfection rates in φ6 bacteriophage are enhanced by virusLinduced changes in host cells. Evol Appl. 2009;2:24–31 10.1111/j.1752-4571.2008.00055.x

39. Hunter M, Fusco D. Superinfection exclusion: A viral strategy with short-term benefits and long-term drawbacks. PLoS Comput Biol. 2022;18:e1010125 10.1371/journal.pcbi.1010125

40. Maffei E, Woischnig A-K, Burkolter MR et al. Phage paride can kill dormant, antibiotic-tolerant cells of *pseudomonas aeruginosa* by direct lytic replication. Nat Commun. 2024;15:175 10.1038/s41467-023-44157-3

41. Lourenço M, Chaffringeon L, Lamy-Besnier Q et al. The spatial heterogeneity of the gut limits predation and fosters coexistence of bacteria and bacteriophages. Cell Host Microbe. 2020;28:390–401. e5 10.1016/j.chom.2020.06.002

42. Shaer Tamar E, Kishony R. Multistep diversification in spatiotemporal bacterial-phage coevolution. Nat Commun. 2022;13:7971 10.1038/s41467-022-35351-w

43. Koskella B, Taylor TB. Multifaceted impacts of bacteriophages in the plant microbiome. Annu Rev Phytopathol. 2018;56:361–80 10.1146/annurev-phyto-080417-045858

44. Simmons EL, Bond MC, Koskella B et al. Biofilm structure promotes coexistence of phage-resistant and phage-susceptible bacteria. mSystems. 2020;5:e00877–19 10.1128/mSystems.00877-19.

45. Bond MC, Vidakovic L, Singh PK et al. Matrix-trapped viruses can prevent invasion of bacterial biofilms by colonizing cells. Elife. 2021;10:e65355 10.7554/eLife.65355

46. Simmons EL, Drescher K, Nadell CD et al. Phage mobility is a core determinant of phage–bacteria coexistence in biofilms. ISME J 2018;12:531–43 10.1038/ismej.2017.190

47. Vidakovic L, Singh PK, Hartmann R et al. Dynamic biofilm architecture confers individual and collective mechanisms of viral protection. Nat Microbiol. 2018;3:26–31 10.1038/s41564-017-0050-1

48. Matange N. Revisiting bacterial cyclic nucleotide phosphodiesterases: Cyclic amp hydrolysis and beyond. FEMS Microbiol Lett. 2015;362:fnv183 10.1093/femsle/fnv183

49. Laganenka L, Sander T, Lagonenko A et al. Quorum sensing and metabolic state of the host control lysogeny-lysis switch of bacteriophage t1. mBio. 2019;10:10.1128/mbio.01884-19 10.1128/mbio.01884-19

50. Lourenço M, Chaffringeon L, Lamy-Besnier Q et al. The gut environment regulates bacterial gene expression which modulates susceptibility to bacteriophage infection. Cell Host Microbe. 2022;30:556–69. e5 10.1016/j.chom.2022.03.014

51. Castledine M, Padfield D, Sierocinski P et al. Parallel evolution of *pseudomonas aeruginosa* phage resistance and virulence loss in response to phage treatment *in vivo* and *in vitro*. Elife. 2022;11 10.7554/eLife.73679

52. Strathdee SA, Hatfull GF, Mutalik VK et al. Phage therapy: From biological mechanisms to future directions. Cell. 2023;186:17–31 10.1016/j.cell.2022.11.017

53. Sanjuán R. The social life of viruses. Annu Rev Virol. 2021;8:183–99 10.1146/annurev-virology-091919-071712

54. Birkholz EA, Morgan CJ, Laughlin TG et al. A mobile intron facilitates interference competition between co-infecting viruses. bioRxiv [Preprint]. 2023:2023.09. 30.560319 10.1101/2023.09.30.560319

55. Isaev AB, Musharova OS, Severinov KV. Microbial arsenal of antiviral defenses–part i. Biochemistry (Mosc). 2021;86:319–37 10.1134/S0006297921030081

56. Trinh JT, Székely T, Shao Q et al. Cell fate decisions emerge as phages cooperate or compete inside their host. Nat Commun. 2017;8:14341 10.1038/ncomms14341

57. Wright RC, Friman V-P, Smith MC et al. Resistance evolution against phage combinations depends on the timing and order of exposure. mBio. 2019;10:10.1128/mbio. 01652-19 10.1128/mBio.01652-19

58. Oz T, Guvenek A, Yildiz S et al. Strength of selection pressure is an important parameter contributing to the complexity of antibiotic resistance evolution. Mol Biol Evol. 2014;31:2387–401 10.1093/molbev/msu191

59. Ferenci T. Irregularities in genetic variation and mutation rates with environmental stresses. Environ Microbiol. 2019;21:3979–88 10.1111/1462-2920.14822

60. Hall AR, De Vos D, Friman V-P et al. Effects of sequential and simultaneous applications of bacteriophages on populations of *pseudomonas aeruginosa in vitro* and in wax moth larvae. Appl Environ Microbiol 2012;78:5646–52 10.1128/AEM.00757-12

61. Weissfuss C, Wienhold S-M, Bürkle M et al. Repetitive exposure to bacteriophage cocktails against *pseudomonas aeruginosa* or *escherichia coli* provokes marginal humoral immunity in naïve mice. Viruses. 2023;15:387 10.3390/v15020387

62. Wienhold S-M, Brack MC, Nouailles G et al. Preclinical assessment of bacteriophage therapy against experimental *acinetobacter baumannii* lung infection. Viruses. 2022;14:33 10.3390/v14010033

63. Gaborieau B, Debarbieux L. The role of the animal host in the management of bacteriophage resistance during phage therapy. Curr Opin Virol. 2023;58:101290 10.1016/j.coviro.2022.101290

64. Van Belleghem JD, Dąbrowska K, Vaneechoutte M et al. Interactions between bacteriophage, bacteria, and the mammalian immune system. Viruses. 2018;11:10 10.3390/v11010010

