## Supplementary Methods for "Phage-phage competition and biofilms reduce the efficacy of a combination of two virulent bacteriophages against *Pseudomonas aeruginosa*"

**Double-agar overlay plaque assay**

For double-agar overlay, 100 µL mid-log suspension of *P. aeruginosa* was poured with 4 mL of Soft Agar (0.7% w/v technical agar BD, 5 mM MgSO_4_ Merck) onto Bottom Agar (1.5% w/v technical agar, BD) plates. Samples and suspensions of interest were 10-fold serially diluted in Phage Buffer (100 mM NaCl, 8 mM MgSO4, 50 mM Tris-HCl, pH 7.5) and were spotted (4 µL/spot) onto the poured soft agar after solidification. Plates were incubated overnight at 37 °C, and 5% CO_2_. The phage titer (are expressed as follows (*n* = number of plaques, *d* = dilution factor, PFU = Plaque forming Unit per mL):

$Phage Enumeration= n*250*d=\frac{PFU}{mL}$.

### **Determination of the frequency of bacteriophage-insensitive mutants**

Bacteriophage-insensitive mutants (BIMs) frequencies were determined as reported earlier [1]. Briefly, a diluted log-culture (OD_600_=0.2) of indicated *P. aeruginosa* and the phage of interest (1E+09 PFU/mL of phage JG005, JG024, or combination of both) was poured into a soft-agar overlay. Number of surviving clones were counted after 24 hours of incubation of plates at 37 °C and 5% CO_2_. The BIMs frequency was calculated as follows:

$BIMs frequency=\frac{number of surviving colonies}{original bacterial titer}$.

**Microtiter biofilm assay: Crystal violet staining**

For biomass determination, the adherent phase was air-dried for 10 minutes after washing. After Crystal violet staining, the content solubilized in 98% ethanol and absorbance was measured at 594 nm (Molecular Devices SpectraMax M2).

**Superinfection exclusion**

PAO1 was infected with either JG005 or JG024 at a high phage-bacterium ratio of 10:1. After 8 minutes of infection, samples were placed on ice for 1 minute and subsequently washed two times with an ice-cold LB medium via centrifugation for 4 minutes at 12.000 xg. Subsequently, the pellet was dissolved with pre-warmed LB medium, and the cells were infected with the respective other phage at a phage-bacterium ratio of 1:10. Phage production was monitored for two hours every 24 minutes.

#### **Bacterial strains and phages used in this study**

**Bacterial strains**

| **Strain name and DSM number** | **Description** | **Source** |
| --- | --- | --- |
| *P. aeruginosa* PAO1  DSM 19880 | Wild-type | DSMZ (Braunschweig, Germany) |
| *P.* *aeruginosa* PA14  DSM 19882 | indicator strain for JG024 phage titration |  |
| *P. aeruginosa* CH3549 [2]^1^  DSM 107571 | Indicator strain for Bhz17 phage titration | TWINCORE, Centre for Clinical and Experimental Infection Research, (Hannover, Germany) and Molecular Bacteriology at Helmholtz Centre for Infection Research (Braunschweig, Germany) |
| *P. aeruginosa* F2230 [2]^2^ | Indicator strain for JG005 phage titration |  |
| *P. aeruginosa* CHA [3] | Bronchopulmonary isolate from patient with Cystic Fibrosis | Jean-Marc Ghigo, Institut Pasteur (Paris, France) |
| *P. aeruginosa* PAO1 Δ*retS* [4] | Deletion mutant of *P. aeruginosa* PAO1 lacking *retS* gene |  |
| ^1^ <https://bactome.helmholtz-hzi.de/cgi-bin/h-disol.cgi?STAT=4&Isol=CH3549> | | |
| ^2^ <https://bactome.helmholtz-hzi.de/cgi-bin/h-disol.cgi?STAT=4&Isol=F2230> | | |

**Pseudomonas lytic phages**

| **Name and**  **DSM number** | **Characteristics** | **Source** |
| --- | --- | --- |
| JG005  DSM 19872 | Class: *Caudoviricetes,* Genus: *Pakpunavirus*,  lytic on *P. aeruginosa* strains PAO1, CHA, F2230 (indicator strain). GenBank: PP712940.1 | DSMZ (Braunschweig, Germany) and Fraunhofer ITEM (Braunschweig, Germany) |
| JG024 [5]  DSM 22045 | Class: *Caudoviricetes,* Genus: *Pbunavirus*,  lytic on *P. aeruginosa* strains PAO1, PA14 (indicator strain).  RefSeq: NC_017674.1 |  |
| Bhz17  DSM 107443 | Non replicating phage on PAO1/PA14 strains. CH3549 is used as the indicator strain |  |
| JG004 [6]  DSM 19871 | Class: *Caudoviricetes,* Genus: *Pakpunavirus*,  lytic on *P. aeruginosa* strains PAO1, CHA, F2230 (indicator strain). RefSeq: NC_019450.1 | DSMZ (Braunschweig, Germany) |
| PTLAW1  DSM 105275 | Phylogenetically similar to JG024 phage (Imke H.E. Korf, unpublished data). Lytic on *P. aeruginosa* strains PAO1, PA14 (indicator strain). |  |

**References**

1.	O'flynn G, Ross R, Fitzgerald G *et al.* Evaluation of a cocktail of three bacteriophages for biocontrol of *escherichia coli* o157: H7. *Appl Environ Microbiol* 2004;**70**:3417-24 <https://doi.org/https://doi.org/10.1128/AEM.70.6.3417-3424.2004>

2.	Hornischer K, Khaledi A, Pohl S *et al.* Bactome—a reference database to explore the sequence-and gene expression-variation landscape of *pseudomonas aeruginosa* clinical isolates. *Nucleic Acids Res*. 2019;**47**:D716-D20 <https://doi.org/https://doi.org/10.1093/nar/gky895>

3.	Dacheux D, Toussaint B, Richard M *et al.* *Pseudomonas aeruginosa* cystic fibrosis isolates induce rapid, type iii secretion-dependent, but exou-independent, oncosis of macrophages and polymorphonuclear neutrophils. *Infect Immun*. 2000;**68**:2916-24 <https://doi.org/https://doi.org/10.1128/iai.68.5.2916-2924.2000>

4.	Goodman AL, Kulasekara B, Rietsch A *et al.* A signaling network reciprocally regulates genes associated with acute infection and chronic persistence in *pseudomonas aeruginosa*. *Dev Cell*. 2004;**7**:745-54 <https://doi.org/https://doi.org/10.1016/j.devcel.2004.08.020>

5.	Garbe J, Wesche A, Bunk B *et al.* Characterization of jg024, a *pseudomonas aeruginosa* pb1-like broad host range phage under simulated infection conditions. *BMC Microbiol*. 2010;**10**:1-10 <https://doi.org/https://doi.org/10.1186/1471-2180-10-301>

6.	Garbe J, Bunk B, Rohde M *et al.* Sequencing and characterization of *pseudomonas aeruginosa* phage jg004. *BMC Microbiol*. 2011;**11**:1-12 <https://doi.org/https://doi.org/10.1186/1471-2180-11-102>
