## Supplemental Table 1 for "Phage-phage competition and biofilms reduce the efficacy of a combination of two virulent bacteriophages against *Pseudomonas aeruginosa*"

**Table S1. List of mutations identified in spontaneous JG005 and JG024 resistance mutant variants of *P. aeruginosa* strains PAO1**

|  | **Clone** | **Susceptibility to** | | | | **Seq ID** | **Effect** | **Locus** | **Gene** | **Gene product** |
| --- | --- | --- | --- | --- | --- | --- | --- | --- | --- | --- |
|  |  | **JG005** | **JG024** | **Meropenem**  **(mm)*** | **Colistin**  **(mm)*** |  |  |  |  |  |
|  | *P. aeruginosa* PAO1 | S | S | 32.2 | 14.5 | CM13 | NA | NA | NA | NA |
| ***P. aeruginosa* PAO1 variants resistant to JG024** | | | | | | | | | | |
| 1 | PAO1 24.1 | S | R/S | 35.7 | 16.8 | CM14 | missense_variant  c.902C>T p.Ala301Val | PA0938 | *wzz2* | hypothetical protein  (modulating O-antigen chain length) |
| 2 | PAO1 24.4 | R | R | 33.9 | 15.9 | CM15 | missense_variant  c.1178C>T p.Pro393Leu | PA3154 | *wzy* | B-band O-antigen polymerase |
| 3 | PAO1 24.6 | S | R | 32.6 | 14.9 | CM16 | stop_gained  c.1006C>T p.Gln336* | PA0938 | *wzz2* | hypothetical protein  (modulating O-antigen chain length) |
| 4 | PAO1 24.8 | R | R | 33.1 | 15.2 | CM17 | frameshift_variant  c.625dupG p.Val209fs | PA3154 | *wzy* | B-band O-antigen polymerase |
| 5 | PAO1 24.10 | R/S | R | 33.4 | 15.1 | CM18 | frameshift_variant  c.136dupA p.Thr46fs | PA3154 | *wzy* | B-band O-antigen polymerase |
| 6 | PAO1 24.12 | S | R/S | 31.4 | 15.2 | CM19 | missense_variant  c.194C>T p.Ala65Val | PA0938 | *wzz2* | hypothetical protein  (modulating O-antigen chain length |
| ***P. aeruginosa* PAO1 variants resistant to both JG005 and JG024** | | | | | | | | | | |
| 7 | PAO1 5-24.3 | R | R | 36.6 | 17.1 | CM21 | frameshift_variant  c.136dupA p.Thr46fs | PA3154 | wzy | B-band O-antigen polymerase |
| 8 | PAO1 5-24.5 | R/S | R | 33.7 | 15.1 | CM22 | frameshift_variant  c.136dupA p.Thr46fs | PA3154 | wzy | B-band O-antigen polymerase |
| ***P. aeruginosa* PAO1 variants resistant to JG005** | | | | | | | | | | |
| 9 | PAO1 5.2 | R/S | R/S | 35.2 | 15.2 | CM24 | missense_variant  c.1214A>G p.Gln405Arg | PA3154 | *wzy* | B-band O-antigen polymerase |
| 10 | PAO1 5.3 | R/S | R | 33.7 | 14.8 | CM25 | frameshift_variant  c.136dupA p.Thr46fs | PA3154 | *wzy* | B-band O-antigen polymerase |
| 11 | PAO1 5.4 | R/S | R/S | 26.8 | 17.9 | CM26 | missense_variant  c.1069G>A p.Gly357Arg | PA2986 | *LolC* | lipoprotein localization protein LolC |
| 12 | PAO1 5.5 | S | R | 32.8 | 15.7 | CM27 | missense_variant  c.538C>T p.Arg180Cys | PA3154 | *wzy* | B-band O-antigen polymerase |
| 13 | PAO1 5.6 | R | R/S | 27.2 | 14.7 | CM28 | missense_variant c.1249T>A p.Trp417Arg | PA5322 | *algC* | phosphomannomutase AlgC  (B-Band O-antigen processing) |
| 14 | PAO1 5.10 | R/S | R | 32.9 | 14.9 | CM29 | stop_gained c.751C>T p.Arg251* | PA3154 | *wzy* | B-band O-antigen polymerase |
| 15 | PAO1 5.1 | R | R | 25.1 | 16.2 | CM23 | ** excluded due to low coverage **; representative clone with brown pigmentation | | | |

**List of mutations identified in spontaneous JG005 and JG024 resistance mutant variants of *P. aeruginosa* strains PA14**

|  | **Clone** | **Susceptibility to** | | | | | **Seq ID** | **Effect** | **Locus** | **Gene** | | **Gene product** |
| --- | --- | --- | --- | --- | --- | --- | --- | --- | --- | --- | --- | --- |
|  |  | **JG005** | **JG024** | **Meropenem**  **(mm)*** | **Colistin**  **(mm)*** | |  |  |  |  |  |  |
|  | *P. aeruginosa* PA14 | NA | S | 33.3 | 15 | | CM30 | NA | NA | NA | | NA |
| ***P. aeruginosa* PA14 variants resistant to JG024** | | | | | | | | | | | | |
| 16 | PA14 SP05 | NA | R | 32 | 17.5 | CM31 | | missense_variant  c.265A>T p.Asn89Tyr | PA14_66120 | | *ssg­* | cell surface-sugar biosynthetic glycosyltransferase, Ssg (small colonies, slower growth) *^1,2^* |
| 17 | PA14 SP14 | NA | R | 32.7 | 14.6 | CM32 | | missense_variant  c.551C>A p.Thr184Asn | PA14_66120 | | *ssg* |  |
| 18 | PA14 SP15 | NA | R | 36.9 | 18.4 | CM33 | | missense_variant  c.551C>A p.Thr184Asn | PA14_66120 | | *ssg* |  |

1. Veeranagouda Y, Lee K, Cho AR, Cho K, Anderson EM, Lam JS. *Ssg*, a putative glycosyltransferase, functions in lipo- and exopolysaccharide biosynthesis and cell surface-related properties in *Pseudomonas alkylphenolia*. FEMS Microbiol Lett. 2011 Feb;315(1):38-45.
2. Wiegand I, Marr AK, Breidenstein EB, Schurek KN, Taylor P, Hancock RE. Mutator genes giving rise to decreased antibiotic susceptibility in *Pseudomonas aeruginosa*. Antimicrob Agents Chemother. 2008 Oct;52(10):3810-3.

**Abbreviation: NA – Not applicable. S – Susceptible; R – Resistant; R/S – reduced susceptibility**

Note 1. Antibiotic susceptibility was determined by a disc diffusion test, in which the meropenem and colistin-impregnated disc (BD BBL™ Sensi-Disc™) was placed on Luria-Bertani agar containing freshly inoculated lawn of individual *P. aeruginosa* strains. Antibiotic and phage susceptibility (see Fig. S5) was determined at least three times. Zone of inhibition is calculated by averaging the three independent readings.

Note 2. Several clones had a second-site polymorphism in an intergenic region (position 4448855, CG->GC) in *P. aeruginosa* PAO1 clones. However, this genomic region is not known to be associated with phages or LPS synthesis and it is also difficult to predict any association with phage resistance or any other functionalities.
